# The time integral of BMP signaling determines fate in a stem cell model for early human development

**DOI:** 10.1101/2023.04.10.536068

**Authors:** Seth Teague, Gillian Primavera, Bohan Chen, Emily Freeburne, Hina Khan, Kyoung Jo, Craig Johnson, Idse Heemskerk

## Abstract

How paracrine signals are interpreted to yield multiple cell fate decisions in a dynamic context during human development *in vivo* and *in vitro* remains poorly understood. Here we report an automated tracking method to follow signaling histories linked to cell fate in large numbers of human pluripotent stem cells (hPSCs). Using an unbiased statistical approach, we discovered that measured BMP signaling history correlates strongly with fate in individual cells. We found that BMP response in hPSCs varies more strongly in the duration of signaling than the level. However, we discovered that both the level and duration of signaling activity control cell fate choices only by changing the time integral of signaling and that duration and level are therefore interchangeable in this context. In a stem cell model for patterning of the human embryo, we showed that signaling histories predict the fate pattern and that the integral model correctly predicts changes in cell fate domains when signaling is perturbed. Using an RNA-seq screen we then found that mechanistically, BMP signaling is integrated by SOX2.

## Introduction

Secreted signaling molecules (morphogens) play key roles in cell fate decisions during embryonic development *in vivo,* as well as in stem cell models *in vitro*^1–5^. However, the relationship between morphogen signaling and cell fate patterning remains incompletely understood. It is generally accepted that the concentrations of signaling molecules determine gene expression and subsequent cell fate choices^6–9^. However, this model does not account for time: concentrations of signaling molecules and downstream signaling activity inevitably change as an embryo develops and cells are therefore unlikely to see a constant signaling environment for the duration of a particular cell fate decision (competence window). This problem is particularly acute in early mammalian development, where no maternal cues are present and signaling gradients are formed in feedback loops by the same cells that differentiate in response to them^4^.

To understand how signaling controls cell fate one should therefore measure the full signaling history rather than focus on a single point in time. This is technically challenging because differentiation takes place in a crowded, ever changing cellular environment and for mammalian cells can take multiple days^10, 11–13^. In addition to the technical challenge, considering signaling histories raises a new conceptual problem. Rather than dealing with a static signaling level as the sole parameter, the signaling history in a cell has a formally infinite number of parameters including rate of signal change, duration, and relative timing of different signals. Therefore, unbiased exploration by direct manipulation of these parameters is impractical, suggesting an indirect approach leveraging spontaneous heterogeneity in signaling activity.

BMP is a key morphogen with a conserved role in dorsoventral patterning across the Bilateria^5, 14^. How BMP controls embryonic patterning has been extensively studied and yet remains controversial. For example, two recent studies in zebrafish came to different conclusions^15, 16^. Current data fall short in at least two respects. First, previous studies do not account for signaling history. Although the BMP signaling gradient in zebrafish and other model systems is known to change over time^15, 17^, it remains unclear how this affects gene expression patterns. In other settings, dynamics of signaling gradients are essential in explaining final gene expression domains^18–20^. Second, studies to date have typically linked average signaling activity with average gene expression. Predicting the approximate fate boundaries along a single axis (such as the dorsal-ventral axis in zebrafish) provides only a few data points with uncertainty introduced by averaging over meaningful heterogeneity such as patterns along the orthogonal axes or subpopulations of cells with qualitatively different signaling dynamics^21^. In contrast, relating signaling to fate in single cells provides thousands of data points in the same embryo, enabling more stringent tests of different models.

Upon BMP4 treatment, hPSCs with precisely controlled colony geometry using substrate micropatterning undergo self-organized spatial patterning into concentric rings of different fates that are specified during gastrulation *in vivo.* This makes micropatterned hPSCs a useful model for human gastrulation known as a 2D gastruloid^22^. Due to its reproducibility and high throughput, this system is ideal for high throughput quantitative studies of differentiation and has led to many insights into the mechanisms of mammalian gastrulation^23–25^, some confirming previous findings in the mouse^26, 27^ and others later confirmed in the mouse^28,29^ or exploring human-specific aspects of development^30^.

Here we used micropatterned hPSCs as a model for early human development to test if and how fate choices in response to BMP might be quantitatively explained by signaling history. To this end we performed live-cell imaging of signaling activity followed by multiplexed immunofluorescence staining to relate signaling to fate in the same cells. We found that combined histories of BMP and Nodal signaling accurately predict cell fate patterns in micropatterned colonies. To limit combinatorial effects of different pathways^27, 30, 31^, we then created conditions to isolate the relationship between BMP signaling and fate. This simplified patterning to a binary decision between epiblast-like and amnion-like cell fates.

To test which features of BMP signaling histories best predicted fate and to establish causality, we complemented analysis of micropatterned hPSCs with experiments in standard culture conditions where increased signaling and cell fate heterogeneity can be leveraged to detect how they are related. We performed automated tracking of signaling related to cell fate in large numbers of individual hPSCs, to our knowledge for the first time. Using an unbiased statistical approach, we showed that measured BMP signaling heterogeneity strongly correlates with cell fate heterogeneity at the single cell level. We found that the initial and final levels of BMP signaling were relatively uniform across cell fates and conditions but that the duration of signaling varied strongly and correlates with cell fate heterogeneity. However, by direct manipulation of signaling level and duration we demonstrated that the level and duration only impact fate by changing the time integral of signaling, which causally determines fate. Thus a lower level of signaling for a longer period of time leads to similar differentiation as higher signaling for a shorter duration, and there is no absolute threshold in the duration or the level of signaling to achieve differentiation.

We then screened for genes that directly reflect the integral of signaling to determine the mechanism by which cells integrate signaling activity over time, which yielded SOX2 as a candidate (among several other genes). We confirmed this using live imaging of endogenous SOX2 and constructed a simple mathematical model that accounts for all of our data by assuming SOX2 represses differentiation genes and decreases in proportion to the time integral of BMP signaling. Finally, we confirmed a prediction of our model in which overexpression of SOX2 would reduce differentiation to amnion-like fate in response to BMP.

## Results

### Signaling dynamics in a stem cell model for human gastrulation predict fate pattern

BMP, Wnt, and Nodal function in a transcriptional hierarchy during self-organized pattern formation in micropatterned hPSCs^26, 27^ (Fig 1A). Wnt and Nodal, as well as FGF signaling are required for primitive streak-like and primordial germ cell-like differentiation, whereas BMP alone is sufficient for amnion-like differentiation^22, 26, 27, 30–32^. We previously measured activity of the BMP and Nodal signaling pathways and found that SMAD4 signaling is dynamic, so static level thresholds cannot account for the final cell fate pattern^33^. Moreover, at the single cell level, BMP signaling at the end of differentiation correlates poorly with cell fate, even under conditions where other signals are pharmacologically inhibited (SI Fig 1AB). Here, we therefore asked if and how dynamic signaling could instead explain the cell fate pattern.

**Figure 1:**
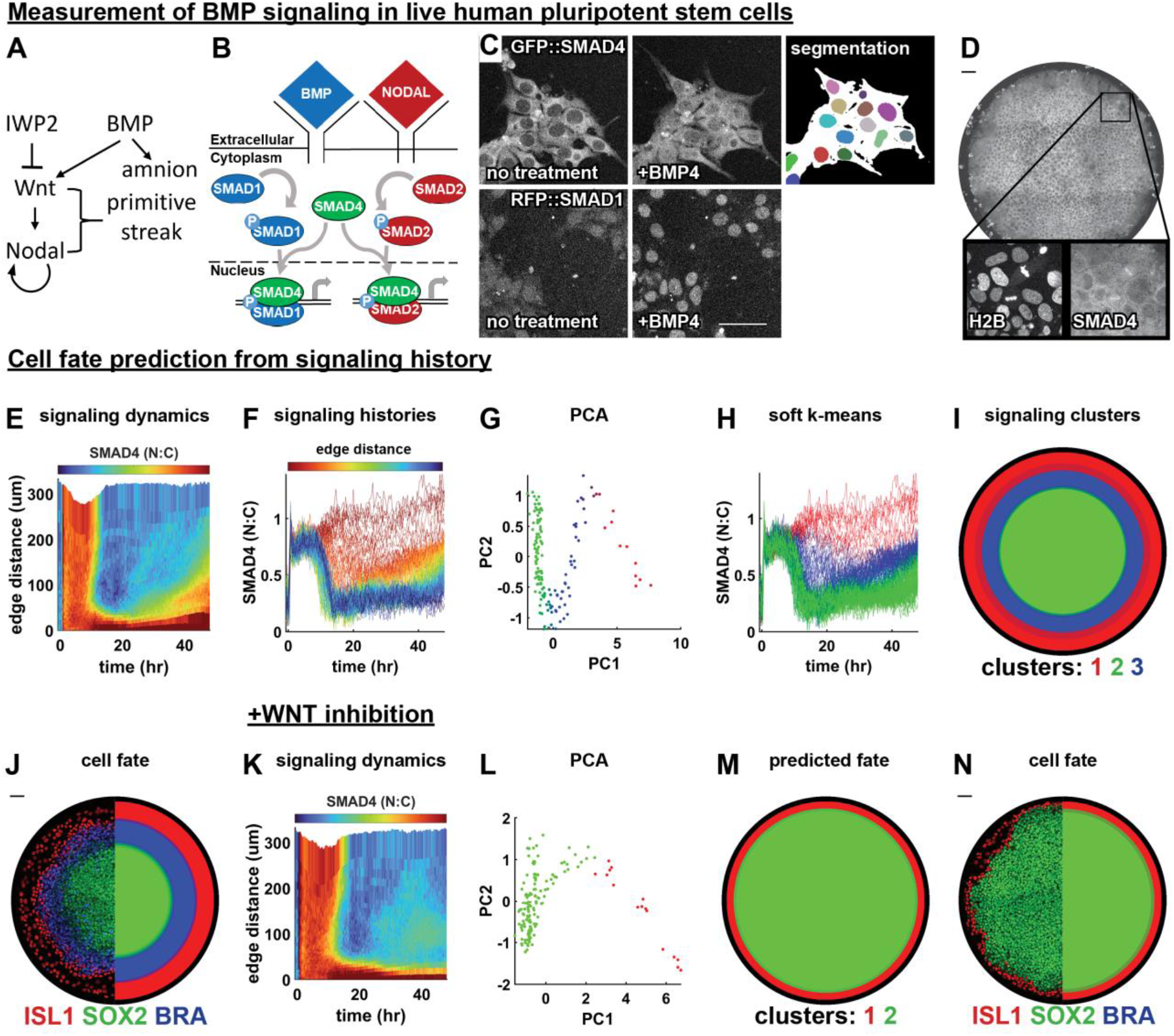
signaling dynamics in a stem cell model for human gastrulation predict fate pattern. (A) Schematic of the BMP, Wnt, and Nodal signaling hierarchy and cell types induced by these signals. (B) SMAD1 conveys BMP signals to the nucleus and SMAD4 conveys both BMP and Nodal signals to the nucleus. (C) Example images showing nuclear translocation of fluorescently tagged SMAD4 (top) and SMAD1 (bottom) proteins in response to BMP4 treatment, and segmentation of nuclei (color) and cell bodies (white) in cells expressing GFP::SMAD4. (D) A representative micropatterned colony of RUES2 cells expressing GFP::SMAD4 at t = 30 hours after treatment with BMP4. (E) A heatmap of average spatiotemporal SMAD4 signaling dynamics (kymograph) in N=5 micropatterned colonies treated with BMP4. (**F**) Plot of radially averaged signaling histories colored for distance from the colony edge. (**G**) Scatterplot of the first two principal components (PCs) of radially averaged signaling histories, colored for soft k means cluster assignment. (**H**) Plot of radially averaged signaling histories colored for cluster assignment. (**I**) Predicted fate map; each radial bin is assigned a color according to the dominant cluster of signaling histories within that bin, over N = 4 replicate colonies. (**J**) Immunofluorescence image of ISL1, SOX2, and BRA in a BMP4-treated colony (left) and the discretized fate map, averaged over replicate colonies (right). Each radial bin is colored for the dominant cell fate within that bin. (**K**) SMAD4 kymograph averaged over N=5 replicate colonies treated with BMP4 and WNTi. (**L**) Scatterplot of the first two PCs of radially averaged signaling histories, colored for cluster assignment. (**M**) Predicted fate map, created as in I. (**N**) IF image of a colony (left) and average fate map (right). Scale bars 50um.

We began by live imaging hPSC colonies over 48h of differentiation with either GFP::SMAD4^34^ or RFP::SMAD1^35^ expression at the endogenous locus and quantified nuclear SMAD levels relative to cytoplasmic levels as a proxy for signaling activity (Fig 1B-D). Although SMAD1 responds only to BMP, SMAD4 responds to both Nodal and BMP (Fig. 1B). To analyze spatiotemporal signaling patterns we exploited the approximate rotational symmetry of the system and averaged signaling over cells at the same distance from the colony edge (Fig 1EF). We reproduced the previously identified pattern in GFP::SMAD4^33^, showing initially uniform signaling that becomes restricted to the edge by 12 hours with a wave of increased signaling beginning around 24 hours. RFP::SMAD1, which had not been measured during patterning before, matched GFP::SMAD4 except for the late signaling wave (SI Fig 1CD), confirming that the late signaling wave reflects Nodal activity, since RFP::SMAD1 does not respond to Nodal^2^.

We then asked if unbiased data analysis could uncover structure in the radially averaged signaling histories. Principal component analysis on the SMAD4 signaling histories revealed a zigzag structure in the signaling histories (Fig 1G) that we computationally divided into 3 clusters (Fig 1H, methods). The cluster means revealed that they represented cells in which signaling was always high (red), high then low (green), or high then low then high again (blue) (SI Fig 1E). When we mapped the signaling clusters back to their spatial positions they formed a spatially coherent pattern even though the clustering did not use spatial information (Fig 1I). We then compared signaling and fate patterns in the same colonies by bleaching fluorescent proteins after live imaging and subsequently staining for fate markers in the same channels (Fig 1J, SI Fig 1G). The resultant fate pattern closely resembled that predicted by signaling (SI Fig. 1I).

We conclude that qualitatively different classes of signaling histories predict cell fate. Our computational approach thereby recovers previous qualitative observations by ourselves and others that sustained BMP signaling leads to amnion-like differentiation while transient BMP followed by Nodal correlates with primitive streak-like differentiation and transient BMP without Nodal remains pluripotent^27, 33, 34^. Nevertheless, this result can be considered surprising for several reasons. First, enough information was provided by measurement of only one signaling protein which provides information about activity of two pathways (BMP, Nodal) out of at least four different pathways that are essential for pattern formation (also Wnt and FGF). Second, this result implies qualitative differences in signaling behavior between the fates: a smooth static signaling gradient would not allow prediction of cell fate domains because it contains no information on where the downstream thresholds determining fate boundaries are. Third, there have been claims that the initial state of a cell predicts its fate^36, 37^ which seems at odds with the signaling determining its fate unless the signaling response and initial state are correlated.

To further challenge our computational approach for predicting fate from signaling, we repeated the analysis after blocking Wnt secretion, which led to the absence of both endogenous Wnt and Nodal signaling due to the Wnt-Nodal hierarchy (Fig 1A). This reduced the pattern to only amnion-like and pluripotent cells. As expected, we no longer observed a Nodal signaling wave (Fig 1K). PCA now yielded only two parts connected by an elbow (Fig 1L), and clustering correctly predicted a binary fate pattern of amnion-like and pluripotent cells (Fig 1MN, SI Fig 1J). The fact that the two signaling clusters are still connected after eliminating the middle cluster from Fig 1G is explained by radial averaging. Eliminating primitive streak-like cells creates a boundary between amnion-like and pluripotent cells, where it leads to averaged signaling between these fates. This suggests that the signaling clusters would be better separated at the single cell level. Overall, these results demonstrate that this novel approach correctly predicts cell fate patterns from signaling and recapitulates known biology even after signaling disruption.

### A pipeline relating single cell signaling history to fate

Having established that qualitatively distinct signaling histories predict the cell fate pattern in micropatterned colonies, we asked if and how specific signaling features control fate. To avoid the full complexity of dynamic combinatorial signaling, we focused on the decision between amnion-like and pluripotent cells controlled by BMP in the absence of WNT (and downstream Nodal). The analysis in Fig. 1 suggests that BMP response is high throughout differentiation in future amnion-like cells. However, it cannot be determined whether there is a minimum level or duration of response required for amnion-like differentiation since only a very small range of levels and durations are represented and each history is an average over a cell population. To address this problem, we therefore developed an experimental and computational pipeline to obtain cell signaling histories linked to fate in individual cells and applied this to a disorganized initial state (i.e. standard culture conditions) to leverage “spontaneous” heterogeneity and sample as broad a range of signaling responses as possible (Fig. 2, methods).

**Figure 2:**
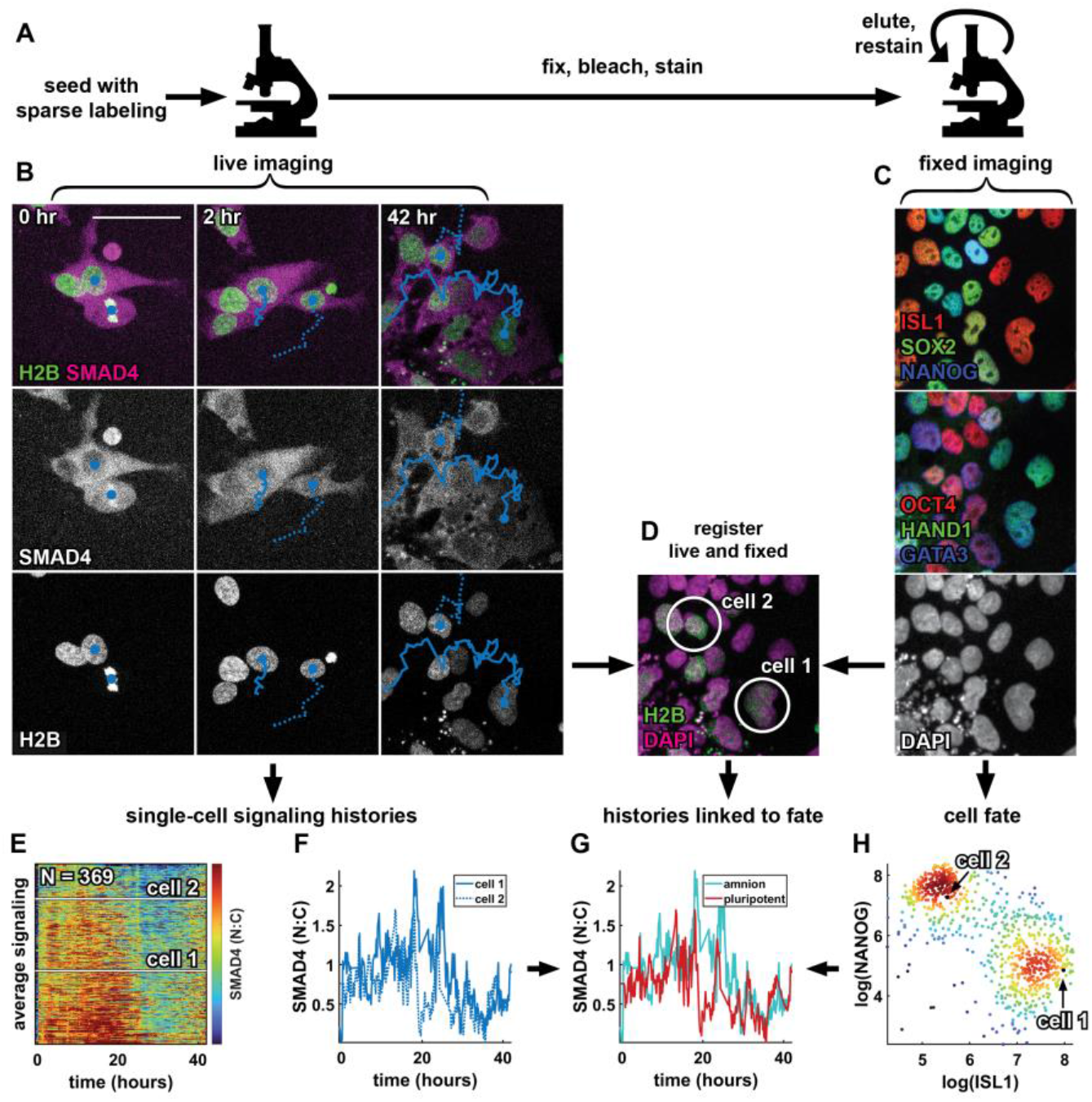
A pipeline relating single cell signaling history to fate. (A) Schematic of the experimental procedure to collect data on signaling histories and cell fate. (B) Example images showing two tracked cells 0, 2, and 42 hours after treatment with BMP4 + WNTi in the SMAD4 and H2B channels. Nuclear centroids are marked with a solid blue circle, tracks are indicated by a solid and dashed line. (C) Multiplexed immunofluorescence data showing an initial stain for ISL1, SOX2, and NANOG (top), followed by a restain for OCT4, HAND1, and GATA3 (middle), in the same field of view as the live data in B. Bottom panel shows the DAPI image from the first round of fixed imaging. (D) Two-color overlay of the live (sparsely labeled) H2B image at 42 hours in B (green) and the DAPI image in C (magenta). The two cells for which tracks are shown in B are circled in white. (E) Heatmap of all single-cell signaling histories collected in a single experiment (N = 369 histories), sorted by mean signaling level. Signaling histories of the two cells tracked in B are marked by white lines. (F) Signaling histories of the two cells tracked in B. (G) Signaling histories of the two cells tracked in B, colored for fate after matching nuclei from the live imaging time lapse to nuclei in the fixed IF data. (H) Scatterplot of log-transformed ISL1 and NANOG intensity in single cells, colored for local density, showing a bimodal distribution of ISL1+ amnion-like cells and NANOG+ pluripotent cells. Scale bar 50um.

To obtain single cell signaling histories we modified a broadly used automated tracking approach^38^ for our conditions, in particular to better handle cell division (SI Fig. 2, methods). Because tracking dense cells over multiple days is challenging we labeled sparsely (10-20%) to obtain more reliable tracks and with manual verification found a linking accuracy of 98.8% between frames. To link signaling histories to fate we fixed cells after live imaging, immunofluorescence stained them and reimaged the same positions. Images of nuclei from live and fixed data were then registered and cells were linked between images to obtain a data structure that contains gene expression data and signaling history for each cell, all in a fully automated manner (Fig 2). Per experiment we could obtain several hundred to a thousand signaling histories linked to fate this way.

Because the cells already express two different fluorescent proteins the number of cell fate markers that could be stained and imaged independently is reduced. As for the micropatterned colonies in Fig.1, we therefore photobleached fluorescent proteins before staining^40^, freeing up all channels. For some experiments we combined this with the 4i multiplexed immunofluorescence protocol to obtain multiple rounds of staining data^41^. Following previous work, we log transformed and normalized expression data for cell fate markers to facilitate downstream analysis (see methods). In summary, we established an automated pipeline to obtain signaling histories corresponding to high-dimensional cell fate data in large numbers of individual hPSCs.

### The time-integral and duration of BMP signaling correlate with cell fate at the single cell level

After creating our experimental pipeline for relating single cell signaling history to fate, we optimized cell density and BMP dose for maximal cell fate heterogeneity, which we expected to be most informative about the relationship between signaling and cell fate. Consistent with previous work^34^ we found that at higher cell densities a higher dose of BMP is needed for a similar level of differentiation (SI Fig 3AB). We decided to initially track GFP::SMAD4 signaling histories for 42h in the medium density condition with an approximately equal split between the fates. After live imaging we then stained for seven transcription factors: ISL1, GATA3, TFAP2C, and HAND1 which mark amnion-like cells and SOX2, NANOG, and OCT4 marking pluripotent cells.

We first analyzed the structure of the signaling histories by themselves. We found individual signaling histories were noisy (Fig. 3A) but their distribution over time had a clear structure and appeared sigmoid, shifting from a relatively steady high level to a low level between 20 and 30 hours (SI Fig. 3C), consistent with previous reports for the mean^33, 34^. To discern the dominant modes of variation between histories we again used principal component analysis. Visual inspection suggested an interpretation for the first three principal components (PCs) corresponding to duration, level of initial response and level of final response (Fig. 3B). To support this interpretation, we directly fitted a sigmoid curve to each signaling history to obtain these features (Fig. 3C) and correlated them with the PCs (SI Fig. 3D). This confirmed PC2,3 respectively showed strongest correlation with high level and low level. However, PC1 correlated strongly with all three signaling features.

**Figure 3:**
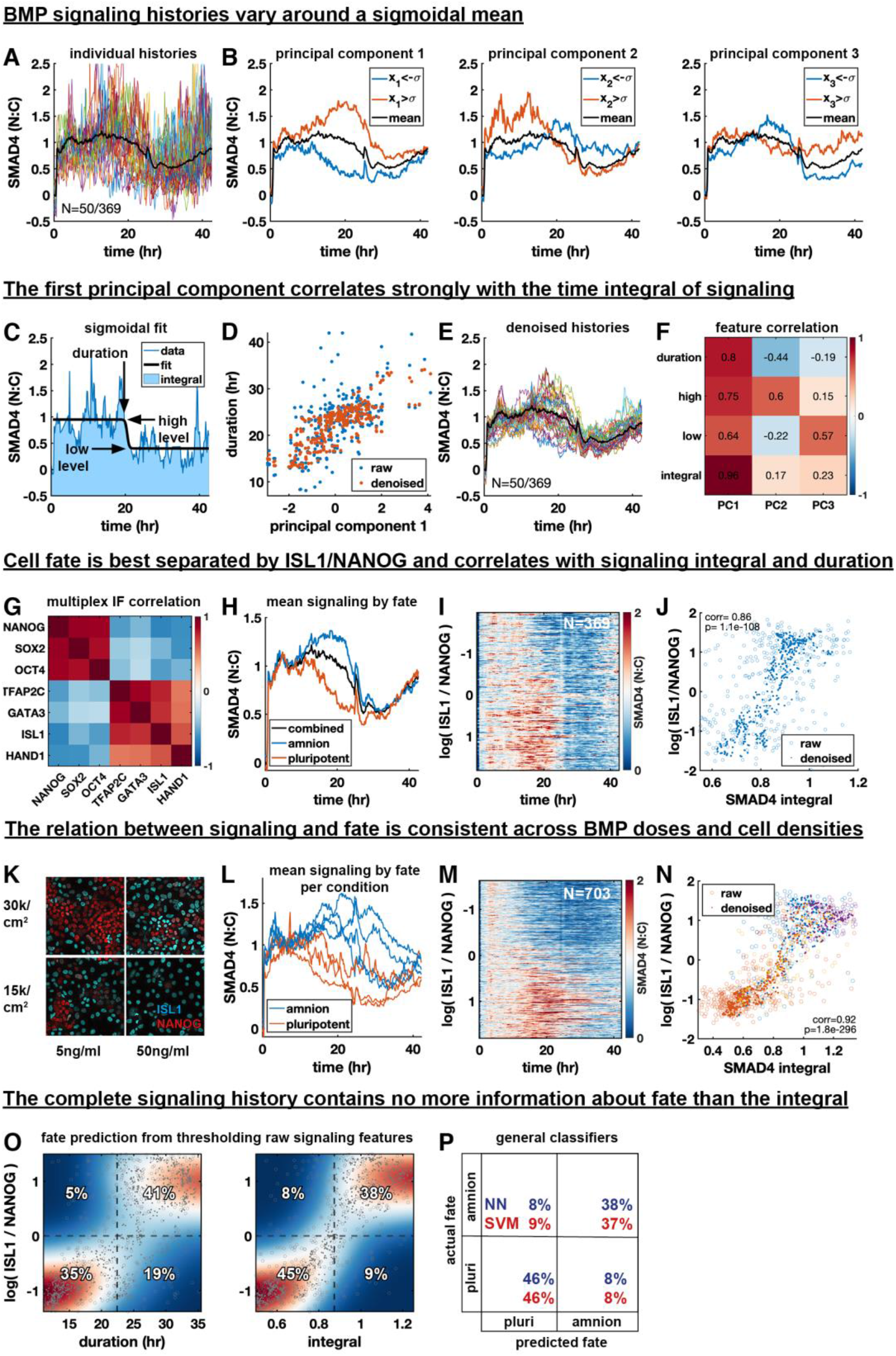
The time-integral and duration of BMP signaling correlate with cell fate at the single cell level. **(A)** Plot of 50 out of 369 signaling histories, population mean overlaid as a bold black line in (A-C). **(B)** Separation of signaling histories along the first three principal components (PCs). Average of histories one standard deviation above or below the mean along each component are shown in red and blue, respectively. **(C)** Example signaling history with a sigmoidal fit to determine signaling features. **(D)** Scatterplot of duration vs. PC1 with and without denoising using MAGIC. **(E)** Signaling histories from (A) after denoising. **(F)** Correlation between signaling features and PCs across cells. **(G)** Correlation of fate marker expression across cells. **(H)** Mean signaling histories for amnion-like and pluripotent cells. **(I)** Heatmap of signaling histories sorted by log(ISL1 / NANOG). **(J)** Scatterplot of log(ISL1 / NANOG) against signaling integral for single cells, with and without denoising. **(K)** antibody stains for ISL1 and NANOG cell densities and BMP4 concentrations. **(L)** Mean signaling for amnion and pluripotent fate for each condition in (N). **(P)** Heatmap of signaling histories sorted by log(ISL1 / NANOG). **(M)** Violin plots of signaling integral distribution for each fate, by condition. **(N)** Scatterplot of log(ISL1 / NANOG) against signaling integral for single cells, with and without denoising. Color is by condition, as in Q. **(O)** Heatmap of kernel density estimate after denoising of conditional distributions of log(ISL1 / NANOG) with respect to duration and signaling integral overlaid with a scatterplots of data points before (circles) and after denoising (dots). Dashed lines show separation of cells into amnion-like and pluripotent based on log(ISL1 / NANOG) or on signaling features. The percentage of cells in each quadrant is indicated, with correct assignments in the top right and bottom left quadrant of each heatmap. **(P)** Confusion matrix showing the performance of a neural network and a support vector machine in classifying cells as amnion-like or pluripotent based on the full signaling history.

In attempting to understand the relationship between PC1 and fitted features we first noticed that the scatterplots of fit parameters versus PC1 contained outliers corresponding to histories with poor fitting of the sigmoid due to noise, to which the duration was most sensitive (Fig. 3D, SI Fig. 3EF). Signaling histories share with single cell RNA-sequencing data that they are noisy high-dimensional single cell measurements. We therefore tested if we could effectively reduce noise in these signaling histories by data diffusion using MAGIC, an algorithm developed for single cell RNA-sequencing data^42^. Weighted averaging of signaling between histories that are most similar using data diffusion visibly reduced noise in signaling histories (Fig. 3E), after which fits improved (SI Fig. 3EF) and outliers disappeared (Fig. 3D), making the relationships between fitted signaling features and principal components more apparent. Although denoising increased the correlation between duration and PC1 relative to other features (Fig. 3F), correlation was still high with all parameters. This suggested that PC1 may represent the time integral (i.e., total amount) of signaling, which increases with any one of the parameters. The resemblance of PC1 to duration in Fig. 3B can be explained by differences in duration dominating variation in the integral. Indeed, we found that the integral of signaling correlates much more strongly with PC1 than other features (Fig. 3F, SI Fig. 3H).

Having discovered duration, high level, low level, and integral as the key features of a signaling history, with integral as the dominant mode of variation, we asked whether these signaling features correlate with cell fate. We first looked at the fate data alone. As expected, the markers separated into two groups representing amnion-like and pluripotent cells (Fig. 3G, SI Fig. 3IJ). Based on the seven-dimensional cell fate data we then clustered the cells into two discrete fates: amnion-like or pluripotent (differentiated or undifferentiated) (SI Fig. 3J-M, methods), and calculated the average signaling history in each cluster. Strikingly, we found that on average the two fates have nearly identical initial and final (high and low) signaling levels but they differ significantly in their mean duration and integral (Fig. 3H). We confirmed this result with RFP::SMAD1 (SI Fig. 3N), although SMAD1 data was much noisier. These key findings show that the dominant mode of variation in BMP signaling, the integral, correlates with the fate.

We then asked if signaling history is not only different between discrete cell fates but whether it predicts expression levels of marker genes, that can be interpreted as coordinates along a differentiation trajectory connecting the fates^43^. We determined ISL1 and NANOG are the respective amnion and pluripotency markers most clearly separating the fates and their difference (after log transforming) log(ISL1) – log(NANOG) = log(ISL1/NANOG) as even better (SI Fig. 3J-M, methods). We therefore focused on this single continuous variable along the fate trajectory for further analysis. A heatmap of signaling histories versus this continuous fate variable showed a clear pattern of higher and longer signaling for more differentiated cells (Fig. 3I). To reveal the relationship between specific signaling features and the degree of differentiation more clearly at the single cell level, we again denoised with controls for artifacts (see methods). Relationships were much easier to discern by eye after denoising. However, there was strong correlation between fate and signaling duration or integral, and less correlation between fate and high or low level (Fig. 3J, SI Fig. 3O) regardless of denoising, and consistent with the means in Fig.3H.

We next asked if the relationship between signaling and fate remains fixed when their distributions are changed by varying cell density and BMP dose. This would be expected in simple models relating signaling and fate, although a more complex context-dependent relationship is possible. We indeed found a consistent relationship between signaling and fate across two different densities and two different BMP doses (Fig. 3K). Mean signaling for amnion-like cells in any condition was separated from mean signaling for pluripotent cells in any condition (Fig. 3L), and the heatmap of combined histories versus fate showed a consistent trend that was much more pronounced than for a single condition (Fig. 3M). Finally, different conditions combined to form a clear threshold-like relationship between signaling features and fate with plateaus for undifferentiated and fully differentiated cells (Fig. 3N, SI Fig. 3P). Importantly, Fig 3L shows that similar to the mean histories of each fate in a single condition (Fig. 3H), the means between conditions show minimal variation in the level of initial response but strong variation in the duration of the response. In other words, different concentrations of BMP lead to different signaling durations but not different signaling levels, consistent with earlier and concurrent work exploring this effect^31, 33^.

Although the integral showed the highest correlation with fate, all features correlated strongly with fate and with each other. We therefore asked if all information about fate is contained in the signaling integral, or whether different features contain independent information. To test this, we first calculated how well an optimal threshold for each feature separates the fates. The percentages of cells in the quadrants formed by the cell fate and signaling feature thresholds in Fig. 3O then provide the confusion matrix of the Bayesian classifier (see methods). The upper right and left quadrants respectively correspond to correctly predicted amnion and amnion misclassified as pluripotent based on the feature threshold, while the lower left and right correspond to correctly predicted pluripotency and pluripotent cells misclassified as amnion. We found the integral-based prediction to be most accurate at 83% (Fig. 3O, SI Fig. 3Q). To estimate the total information contained in signaling we trained a variety of general classifiers, specifically artificial neural networks and support vector machines, and found their accuracies in predicting fate from the complete signaling history to be extremely similar, also around 83% (Fig. 3P). This suggests that the complete signaling history contains no more information about fate than the integral. Although we conservatively performed this analysis with raw data to exclude the possibility of denoising artifacts, we found that after denoising, integral-based prediction becomes nearly perfect (97% accuracy), further supporting the conclusion that all information about fate is contained in the integral.

In summary, we found that the signaling response to BMP is characterized by a high initial plateau going down to a lower final plateau. Although it is primarily the duration that varies between cell fates and experimental conditions, fate is best explained by the time integral of signaling which in turn depends on the duration. This raises the question of whether mechanistically either the time-integral or the signaling duration controls fate.

### The time-integral of BMP signaling controls differentiation

Having found that both the duration and time-integral of signaling strongly correlate with fate, we asked how integral-dependent differentiation can be distinguished from a level threshold combined with a duration threshold, which is the simplest dynamic extension of the classic morphogen model. We reasoned that if the integral controls cell fate, lower levels of signaling can be compensated by a longer duration, which is inconsistent with an absolute level threshold (Fig. 4AB).

To independently control signaling level and duration and determine how their combination impacts fate, we could not use treatment with different concentrations of BMP4, since this primarily changes the duration of response^31, 33^ (Fig. 3L). Therefore, we first combined low cell density with high BMP concentration to obtain a high BMP response in all cells throughout differentiation. We then controlled the signaling level by treatment with different doses of the BMP receptor inhibitor LDN193189 (BMPRi) and the duration by BMP removal combined with a high dose of BMPRi to abruptly and completely shut down signaling (Fig. 4C, SI Fig. 4A). Accordingly, the signaling level was experimentally defined as the mean signaling before signaling shutdown and the duration as the time of the shutdown.

**Figure 4:**
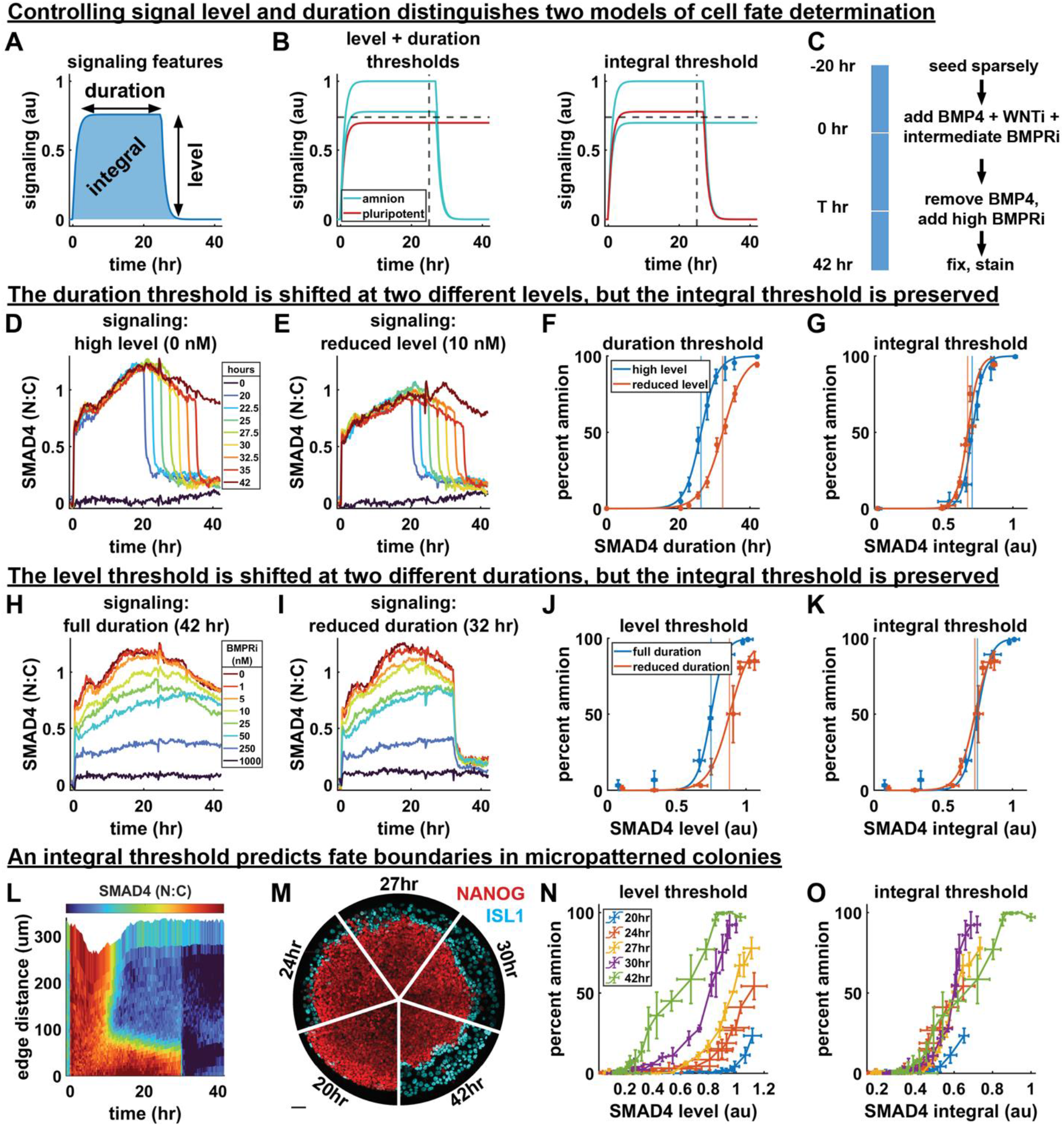
The time-integral of BMP signaling controls differentiation. (**A**) Diagram of relevant signaling history features. (**B**) Hypothetical set of signaling histories for which a combined level and duration threshold model makes a different prediction of cell fate than an integral threshold model. (**C**) Schematic of the experimental procedure used to control the level and duration of signaling. (**D**) Mean signaling for different durations without initial BMPRi. (**E**) Mean signaling for different duration with an initial BMPRi treatment of 10 nM, legend in 4D. (**F**) Duration thresholds based on logistic sigmoid fit for two signaling levels, error bars are standard deviation over N = 4 technical replicates. Thresholds in any signaling feature are defined by 50% differentiation. (**G**) Integral thresholds for the same data as (F). (**H**) Mean signaling for 8 doses of BMP inhibition with LDN193189. (**I**) Mean signaling for different initial levels of BMPRi with a 32-hour duration. (**J**) Level threshold for both signaling durations. Error bars are standard deviation over N = 3 technical replicates (**K**) Integral thresholds for data from (J). (**L**) Kymograph of average BMP signaling for N=3 colonies treated with BMP inhibitor at 30 hours. (**M**) IF data showing amnion differentiation for each signaling duration (scale bar 50um) (**N**) Percent differentiation against mean signaling level before shutdown for each duration. Each point represents a radial bin (a fixed distance from the colony edge). Error bars are standard deviation N=3 colonies per condition. (**O**) Percent differentiation against signaling integral. Error bars as in (N).

We first varied duration while holding level fixed. Consistent with our findings in Fig. 3 we found a sharp duration threshold around 26 hours, above which cells predominantly differentiated to amnion-like cells (Fig. 4D, SI Fig. 4B). We then repeated this while lowering the level with a small dose of BMPRi throughout the experiment (Fig. 4E, SI Fig. 4C) and found that the duration threshold went up to around 32 hours (Fig. 4F), consistent with our hypothesis that there are no fixed level and duration thresholds. Moreover, when we plotted the duration-differentiation curves for two different levels (Fig. 4F) against the time-integral of signaling, these collapsed on top of each other with an identical integral threshold (Fig. 4G). Of note, the duration threshold in Fig. 4D is slightly later than that found in Fig. 3, which is also consistent with integral control of fate because the final signaling level after BMPRi treatment is lower than in cells where the BMP response goes down spontaneously. Overall, the fact that the duration required for differentiation changes at different signaling levels but the integrated signaling remains the same provides strong quantitative support for the integral model.

To further support our integral hypothesis, we then explored a wide range of signaling levels while holding the duration fixed. We found a sharp threshold in level above which pluripotency was lost (Fig. 4H, SI Fig. 4D) that was shifted upward when the duration of signaling was decreased (Fig. 4IJ, SI Fig. 4E). These level-differentiation curves collapsed on a common integral threshold (Fig. 4K), again quantitatively supporting the idea that the integral controls fate and ruling out an absolute threshold in signaling level.

Finally, we asked if the integral model can also account for differentiation in the micropatterned model for embryonic patterning, where a BMP signaling gradient forms spontaneously from the edge inward due to receptor accessibility and secretion of inhibitors^28, 30^ (SI Fig. 4F). If BMP signaling is shut down across the colony, cells at different distances from the edge will have been at different signaling levels for the same duration and therefore have experienced different amounts of integrated signaling (SI Fig. 4G). By shutting down BMP signaling in micropatterned colonies at different times and measuring differentiation as a function of distance from the colony edge we are then simultaneously testing the effect of level and duration. We therefore live-imaged micropatterned colonies of GFP::SMAD4 hPSCs, treated them with a high dose of BMPRi at different times and then fixed them after 42h to evaluate the percentage of amnion-like cells at different distances from the edge (Fig. 4LM, SI Fig. 4H-P). Consistent with the experiments in sparse culture, we found that the signaling level required for 50% differentiation was strongly dependent on the duration of signaling (Fig. 4N, SI Fig. 4Q), but that radius-differentiation curves approximately collapsed on a common integral threshold (Fig. 4O).

It has been claimed that GATA3 acts as an irreversible switch driving commitment to differentiation after as little as one hour of BMP signaling, which seems inconsistent with our findings. For direct comparison we therefore measured GATA3 in the same experiment (SI. Fig 4R). We found that similar to ISL1 it reflects the integrated signaling, but surprisingly does not show a switch-like threshold, appearing graded instead (SI Fig. 4ST). We therefore found no evidence of a BMP signaling duration threshold for GATA3 activation, early or late. Altogether, our data provide strong evidence that amnion-like differentiation is controlled by the time-integral of BMP signaling across in both standard culture and micropatterned colonies and that there are no absolute thresholds in level of duration or level of signaling.

### BMP signaling is integrated by SOX2

We asked by what mechanism cells integrate BMP signaling and reasoned that the simplest mechanism would be a protein that increases or decreases at a rate that is proportional to the level of BMP signaling on a timescale comparable to that of differentiation. A threshold response of differentiation markers like ISL1 or HAND1 downstream of such an integrator gene would then explain the integral threshold observed in Fig. 4. We therefore looked for genes showing gradual increase or decrease on the timescale of differentiation with immediate response at a rate roughly proportional to the level of BMP signaling.

First, we measured protein-level dynamics of amnion and pluripotency genes for different levels of BMP signaling using immunofluorescence staining on a timeseries of fixed samples. We observed three classes of dynamics (Fig. 5A,D, SI Fig. 5A): gradual increase with a dose-dependent slope (GATA3, TFAP2C), gradual decrease with a dose-dependent slope (SOX2, NANOG), and delayed increase (ISL1, HAND1). For genes showing immediate response we related their rate of change to the level of SMAD4 signaling for the same dose of BMPRi (Fig. 5B, SI Fig. 5B) in the first 24h and found an approximately linear relationship for each (Fig. 5C, SI Fig. 5C). In contrast, for ISL1, HAND1, and OCT4 we confirmed the threshold dependence on SMAD4 of Fig. 4 (SI Fig. 4D). Therefore this analysis identified four of these seven genes as potential integrators (GATA3, TFAP2C, SOX2, NANOG).

**Figure 5:**
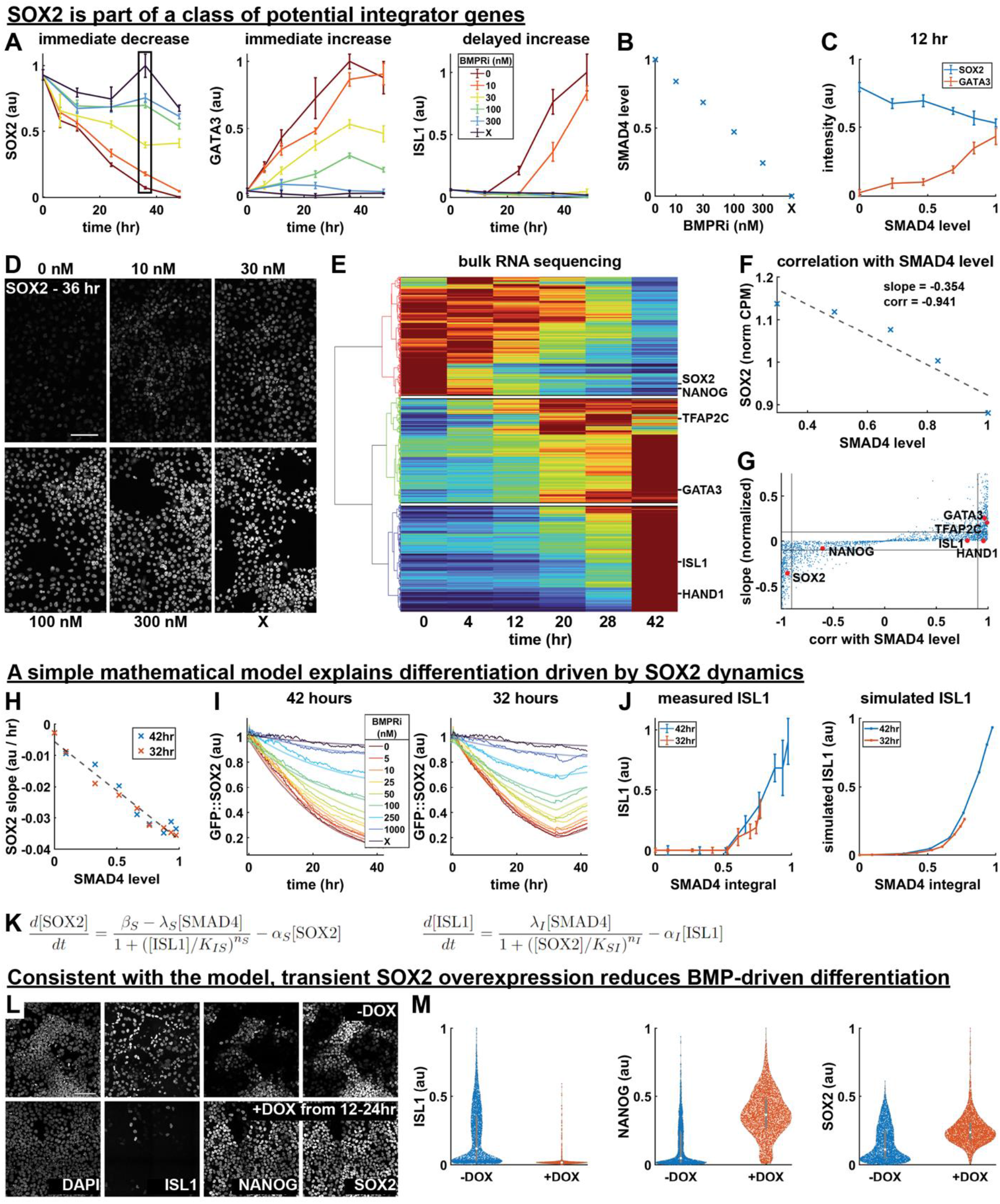
BMP signaling is integrated by SOX2. (**A**) Normalized expression of SOX2, GATA3, and ISL1 over time for different signaling levels, measured with time-series IF. Error bars are standard deviation across N = 6 images. A box outlined in black shows the data points corresponding to image data in 5D. (**B**) Average SMAD4 level for each treatment condition in 5A, determined using SMAD4 dynamics measured in the same conditions and shown in SI fig 5B. (**C**) SOX2 and GATA3 expression at 12 hours, plotted against SMAD4 signaling level. (**D**) Example IF image data for SOX2 in each treatment condition at 36 hours, corresponding to the points boxed in 5A. (**E**) Heatmap of time-series bulk RNA seq data (normalized counts per million) with genes on the y-axis ordered by hierarchical clustering. The cluster dendrogram is shown to the left, with lines colored for discrete cluster assignment, and white lines are drawn on the heatmap to separate clusters. The location in the heatmap of genes also measured with IF is indicated to the right. (**F**) Example bulk RNA seq dose-response data is shown for SOX2 with a linear least squares fit of SOX2 with respect to SMAD4 level. The slope of the least squares fit and the correlation coefficient between SOX2 and SMAD4 level are indicated. (**G**) Scatterplot of the slope of each gene with respect to SMAD4 and that gene’s correlation with SMAD4 in the dose-response bulk RNA seq data, as determined in 5F. The locations in the scatterplot of genes for which we also have IF data are indicated. (**H**) Plot of the slope of each GFP::SOX2 curve in SI fig 5G over the first 16 hours of differentiation against SMAD4 signaling level. SMAD4 level is determined in each condition based on data in Fig 4D. A dashed line indicates a linear fit of the data. (**I**) Measured (thick, solid lines) and simulated (thin, semi-transparent lines) GFP::SOX2 dynamics over the course of 42 hours of differentiation with indicated treatments applied for 42 (left) or 32 (right) hours. (**J**) Measured (left) and simulated (right) ISL1 level as a function of SMAD4 integral for the conditions in 5J. (**K**) Equations used to model the regulation of SOX2 and ISL1 by BMP-SMAD4 signaling, and their mutual inhibition. (**L**) Example IF data showing ISL1, SOX2, and NANOG expression after 42 hours of differentiation with 50 ng/mL BMP4 + WNTi, with or without the addition of doxycycline from 12 to 24 hours. (**M**) Violin plots of measured ISL1, NANOG, and SOX2 expression levels in treatment conditions in 5L. All scale bars 100um.

We then performed similar analysis at the transcriptional level in a genome-wide, unbiased manner. We screened for integrator genes with bulk RNA-seq at different times after BMP treatment and at different levels of BMP signaling after 5h. To focus on genes with large expression changes we restricted our analysis to genes with a cumulative fold change over time of at least one standard deviation above the mean (SI Fig. 5E). Consistent with the immunofluorescence data, hierarchical clustering of the time series then revealed three clusters of genes undergoing large changes in expression during differentiation: 818 continuously decreasing genes including SOX2 and NANOG, 734 immediately increasing genes containing GATA3 and TFAP2C, and 730 delayed increasing genes including HAND1 and ISL1 (Fig. 5E, SI Fig 5F). Other amnion markers^44, 45^ were in the increasing clusters: TFAP2A and GATA2 were early response genes and GABRP, KRT7, TP63, and CDX2 were late increasing genes (SI Fig 5F). We then identified genes with an immediate response to BMP that was both proportional and strong, by respectively filtering based on the correlation with SMAD4 level (above 0.9) and slope (above 0.1 normalized to the time-series maximum) in the dose-response data after 5h (Fig 5FG). Intersecting these with the set of genes from the timeseries analysis left 264 immediately decreasing and 244 immediately increasing genes as candidate integrators. Reassuringly, these respectively contained SOX2 and GATA3, TFAP2C. However, NANOG was excluded due to low correlation with SMAD4 signaling at 4h. This suggests NANOG is not a direct transcriptional target of BMP signaling and its response on longer timescales measured with IF is either indirect or post-transcriptional. Overall, the bulk RNA-seq data identified potential integrator genes and provides a deep characterization of the transcriptional response to BMP4 in hPSCs.

Decreasing genes are associated with the pluripotent state and increasing genes with amnion-like fate, raising the question of whether the integral threshold represents a loss of pluripotency or a commitment to amnion fate. These are indistinguishable in our experiments because we excluded other fates with Wnt inhibition. However, we found that ISL1 and HAND1 anticorrelate more strongly with SOX2 and NANOG than they correlate with GATA3 and TFAP2C (Fig. 3G), suggesting that loss of pluripotency genes may be more important than gain of expression of early amnion genes to drive expression of late amnion genes. Furthermore, it was recently proposed that there is a time window in which cells expressing amnion markers can still acquire primitive streak-like fate by exposure Wnt^31^. This also suggests that our integral threshold represents a commitment to differentiate, i.e. loss of pluripotency, rather than commitment to amnion-like fate. To single out a specific candidate integrator gene we therefore decided to focus on pluripotency genes. We then identified SOX2 as the most likely candidate since it was the only of the so-called ‘core pluripotency genes’ that fit the criteria for an integrator in the bulk RNA-seq analysis.

To test how well SOX2 levels reflect integrated SMAD4 signaling, we again varied levels of BMP signaling for two different durations as in Fig. 4, but now measured SOX2 over time using a cell line expressing GFP::SOX2 in the endogenous locus (Fig 5I). With SMAD4 signaling inferred from GFP::SMAD4 under the same conditions (Fig 4, SI Fig 5G), GFP::SOX2 dynamics during the first 16 hours of differentiation confirmed the linear relationship between SMAD4 level and the rate of SOX2 decay over a wide range of SMAD4 levels, as expected if SOX2 integrates SMAD4 signaling (Fig 5HI) and consistent with similar observations by Camacho-Aguilar et al.^31^

Our data also showed deviations from the linear relationship at later times, in particular recovery after shutdown of BMP signaling, likely due to protein turnover. We asked whether a simple model with production and degradation of SOX2 leading to exponential time-dependence could explain these deviations and still be consistent with our measured integral threshold. To answer this question we implemented this model mathematically. We included ISL1 downstream of SOX2 with a threshold dependence (SI Fig. 5H) which resulted in SOX2 levels reflecting a weighted integral of BMP signaling that approximates the true integral for slow enough turnover (model SI). The simple exponential model showed generally good agreement with both the observed SOX2 dynamics and the measured relationship between ISL1 expression and integrated BMP signaling (SI Fig. 5H). However, for high levels of BMP signaling, SOX2 did not plateau as expected and average SOX2 recovery rates were lower than expected after BMP shutdown at 32h. We reasoned that this was due to differentiation: SOX2 becomes permanently repressed and therefore does not recover in differentiated cells, which constitute a larger fraction after exposure to higher levels of BMP signaling. We modeled this at the level of the population means by adding negative regulation of SOX2 by ISL1, which resolved the observed discrepancies (Fig. 5IJK).

To test the role of SOX2 directly we then created a doxycycline inducible SOX2 cell line (SI Fig 5IJ). We found that doxycycline-induced SOX2 overexpression for all 42h of differentiation prevented upregulation of amnion markers, suggesting SOX2 represses these genes (SI Fig 5KL). However, NANOG expression was also lost, consistent with the known requirement for the right stoichiometry between pluripotency genes to maintain pluripotency^46, 47^. To stay within the range of SOX2 levels where pluripotency is possible, we then treated cells with doxycycline for 12h after 12h of BMP treatment, when endogenous SOX2 levels have already decreased significantly. We found that this significantly decreased differentiation to ISL1+ amnion but maintained pluripotency in the ISL1-negative cells, directly demonstrating that SOX2 level controls the differentiation threshold (Fig 5LM).

## Discussion

We showed that the time-integral of BMP signaling determines amnion-like fate in human pluripotent stem cells. We also identified ‘integrator genes’, whose levels reflect the time integral of BMP and provide evidence that SOX2 mechanistically implements the time-integration of BMP signaling. Importantly, this challenges the idea of level thresholds controlling differentiation since the same integrated signaling level can be reached by short high signaling and long low signaling. These findings therefore potentially have broad repercussions for our understanding of developmental patterning, while providing specific insight into early human cell fate decisions and heterogeneous stem cell differentiation *in vitro*. We investigated how signaling is interpreted by cells but did not ask how signaling heterogeneity arises. Previous work suggests this may be due to receptor localization and inhibitor secretion combined with local differences in cell density^28^.

In interpreting our data, we made several idealizations. First, for integrator genes to perfectly reflect the integral of BMP signaling they would have to be infinitely stable, whereas real integrator proteins are expected to have finite lifetime. Our mathematical model showed that this is a good approximation, although larger deviations from an exact integral threshold are predicted for low levels combined with longer durations than we experimentally tested. Second, we used the term fate for ISL1+ amnion-like cells versus SOX2+ pluripotent cells, but as this is the first step of differentiation from pluripotency in a long series of developmental events, we do not expect the differentiation markers to truly mark a stable cell fate. Rather, our amnion-like cells may represent an intermediate state towards further differentiation. Nevertheless, we showed commitment to differentiation, as the pluripotency gene SOX2 does not recover even after removal of BMP in cells that pass the threshold for differentiation (Fig 5).

There are several potential advantages to integrating cell signaling over time. Integration reduces noise and is insensitive to brief signaling perturbations in the same way as averaging. It also allows for flexible tuning of fate patterns, since an integral threshold is equivalent to a level threshold that can be tuned by changing the duration, e.g. with a delayed negative feedback loop, which may be more straightforward than scaling the morphogen gradient itself. This also implies that if the duration of signaling is the same for all cells, a signaling gradient interpreted by integration over time would produce cell fate patterning consistent with the classic French flag model despite the absence of absolute thresholds in signaling level.

Given these advantages, temporal integration of cell signals might be expected throughout development and indeed there is evidence for integration of ERK signaling^48^. In many other contexts where a strong dependence on signal duration was found, this could be a dependence on integrated signaling, depending on whether duration thresholds change with signaling level. For example, our finding that BMP dose primarily affects the average duration of signaling (Fig. 3) is reminiscent of SHH signaling in the neural tube, where concentration increases increase the duration but not the level of signaling^18, 20^. Strikingly, BMP signal duration is also important in neural tube patterning, but it remains unclear whether dorsal interneuron identity depends on the time-integral of BMP signaling or on level, duration, and other signaling features separately^17^. In addition to the time-integral of signaling possibly controlling fate more generally, the mechanism of BMP integration by SOX2 may also extend to other cell fate decisions, since several fate decisions controlled by BMP involve suppression of SOX2 including neural versus non-neural ectoderm^49–51^ and foregut versus hindgut^52^. An important related question is whether the sigmoidal signaling dynamics we found in response to BMP are typical or depend on developmental stage and the combination of BMP ligands and receptors^53, 54^.

Several papers previously considered the role of BMP signaling in hPSC differentiation. Gunne-Braden et al.^55^ claimed that GATA3 mediates fast, irreversible commitment to differentiation after less than 1h of BMP exposure. In sharp contrast, our work shows that a substantial duration (over 24h) of BMP signaling is required for both differentiation and high GATA3 expression even at maximal signaling levels. Moreover, SI Fig. 4I-K and Fig. 5 show a gradual relationship between GATA3 expression and the duration of BMP signaling that supports our integral model and is inconsistent with switch-like behavior. Consistent with our data, a large body of literature supports the conclusion that BMP inhibition at any time during differentiation has a clear impact on cell fate^27, 28, 34, 56–58^.

Tewary et al.^32^ also studied how BMP concentration and duration affect micropatterned hPSC colonies. However, they only mathematically modeled BMP gradient formation and did not quantitatively investigate the relationship between signaling and fate. Qualitatively, they proposed final pSmad1 levels determine fate boundaries but this seems inconsistent with their finding that marker genes are expressed earlier at higher doses of BMP. BMP signaling in micropatterned colonies does not increase over time (Fig. 1) so cells above a pSmad1 level threshold for a given marker would be expected to be above it at all times and thus show similar expression dynamics. In contrast, the integral model predicts earlier expression at higher signaling levels. Their data therefore appear consistent with our integral model.

Nemashkalo et al.^34^ showed duration rather than the initial BMP response correlates with fate at the population level and proposed a duration threshold. Although they performed single cell tracking, this was for a small number of cells in a single condition and not for the full duration of differentiation. Consequently, they were unable to quantitively relate signaling to fate and did not demonstrate a fixed duration threshold. They also did not study large micropatterned colonies that model embryonic patterning. In contrast, here we showed there is no duration threshold but rather an integral threshold and we provide a downstream mechanism that quantitatively explains fate from signaling in both standard culture and micropatterned colonies.

Finally, concurrently with and complementary to this work, Camacho-Aguilar et al.^31^ explored the effect of concentration versus duration of BMP treatment on cell fate in a different context where BMP acts combinatorially with downstream Wnt to control a decision between primitive streak-like and amnion-like fate. In contrast, we were able to determine which features of BMP signaling history control gene expression only by inhibiting Wnt. However, their results are consistent with ours in showing that different concentrations of BMP primarily affect the duration of response.

Future work will have to address several details regarding the molecular mechanism of BMP integration. For example how the rate of SOX2 decrease is controlled by BMP signaling and how SOX2 represses late differentiation genes including ISL1. Perhaps most importantly, we do not understand why there are early differentiation genes including TFAP2C and GATA3 which also meet all the requirements to be integrator genes in addition to late differentiation genes such as ISL1 and HAND1 whose expression appears to mark a commitment to differentiation. One possibility is that these genes act as integrators complementary to SOX2, e.g. our data are consistent with the possibility that ISL1 upregulation requires both SOX2 downregulation and GATA3 upregulation.

There has been debate about whether heterogeneous differentiation in human pluripotent stem cells is primarily due to a heterogeneous initial state or heterogeneous signaling response^59^ and several papers found initial levels of pluripotency markers to be predictive of differentiation^36, 37^ although the simplest models for tissue patterning assume a fixed relationship between signaling and fate. Intuitively it is clear that both should matter and that it depends on the specific context which dominates. However, our findings unify these contrasting results in the literature more concretely by providing a direct connection between the initial levels of the pluripotency factor SOX2, which sets the threshold for integrated BMP signaling, allowing calculation of the relative contributions of signaling heterogeneity and initial heterogeneity in SOX2.

Our tracking approach in standard culture is easily generalized to widely used differentiation protocols and future work will investigate whether the same mechanism for BMP interpretation is reused at different developmental stages. Equally important is investigating at the single cell level how combinatorial signaling histories are interpreted, for example in the human germline, where the relative timing and duration and BMP and Nodal signaling appear to play a key role in determining fate.

## Acknowledgements

We thank Mara Duncan, Ben Allen, Doug Engel, and Aryeh Warmflash for discussions and feedback on the manuscript. This work was supported by the National Institute of General Medical Sciences (NIGMS R35GM138346), NIH Cellular Biotechnology Training Program (CBTP) T32GM145304, the Branco Weiss Fellowship – Society in Science, and the University of Michigan.

## Methods

### Cell lines

We used the embryonic stem cell lines ESI017 (XX) and RUES2 (XX; Ali Brivanlou, Rockefeller). Genetically modified variants of these two cell lines were RUES2 GFP::SMAD4 (Nemashkalo et al., 2017), RUES2 RFP::SMAD1 (Yoney et al., 2018), and ESI017 tetO-SOX2 (this paper). We additionally used the genetically modified induced pluripotent stem cell line WTC11 GFP::SOX2 (XY; Allen Institute, identifier AICS-74; RFP nuclear marker added by us).

### Cell culture and differentiation

Human pluripotent stem cells were cultured in the pluripotency-maintenance media mTeSR1 (StemCell Technologies) on Cultrex (R&D Systems)-coated 35mm tissue culture plates. During routine maintenance, cells were passaged every 2-4 days, either in whole colonies with L7 (Nie et al., 2014), or in single-cell suspension with Accutase. Seeding for differentiation experiments was done in single-cell suspension after dissociation with Accutase. When passaging in single-cell suspension with Accutase, cells were kept for 24 hours in ROCK inhibitor (RI) after passaging. RUES2 GFP::SMAD4 cells were selected with 24 hours of blasticidin treatment at every passage. Blasticidin was removed 24 hours after passaging, and was always excluded during differentiation experiments.

For differentiation experiments in standard (not micropatterned) culture, cells were passaged and seeded into μ-Slide 18-well plates from Ibidi (with the exception of bulk RNA seq experiments, which used 24-well Ibidi plates) 16-20 hours before treatment with BMP. When performing live-cell imaging, media was changed from standard mTeSR to mTeSR without phenol red 2.5 hours before BMP4 treatment. For experiments in which cells were not maintained in RI over the course of differentiation, RI was removed 2.5 hours before treatment time. For experiments in figure 3, cells were sparsely labeled to facilitate automated single-cell tracking. For sparse labeling, 10-20% GFP::SMAD4 RUES2 cells were mixed thoroughly with 80-90% RUES2 parent cells and seeded at the desired density.

To differentiate cells in micropatterned colonies, we followed the procedure described in Jo et al., 2022^30^, adapted from the protocol in Deglincerti et al., 2016^60^. Briefly, cells were dissociated with Accutase, resuspended in a single-cell suspension, and seeded at 470k cells/cm^2^ onto laminin-coated micropatterns in mTeSR with RI. Colonies were washed 2x with PBS-/- 45 minutes after seeding to clear away those binding non-specifically in the well outside of micropatterned colonies. Two hours after seeding, RI was removed and BMP4 treatment added. Micropatterning experiments were performed in 18-well Ibidi slides prepared according to the protocol in Azioune et al., 2009^61^. All micropatterned colonies had a diameter of 700 μm. Signaling reagents and treatment concentrations are listed in Table 1.

**Table 1:**
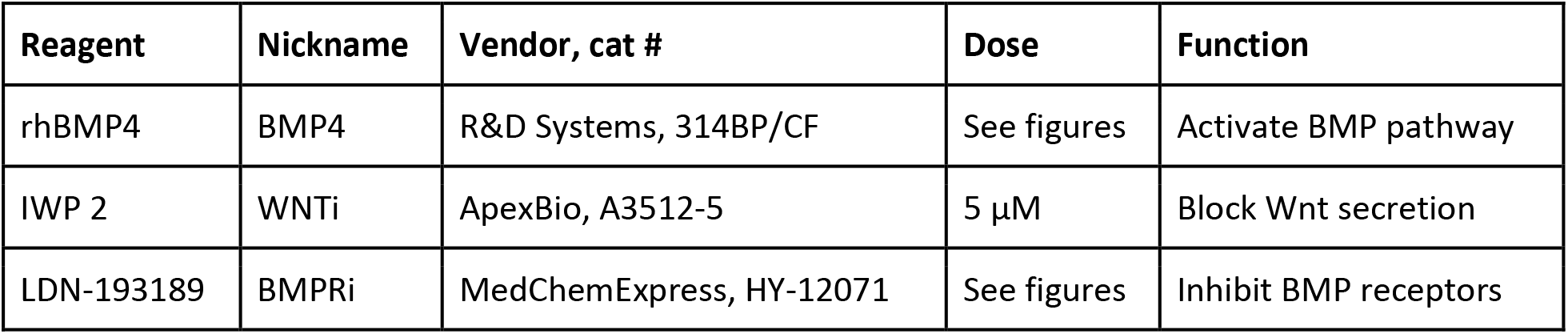
Cell signaling reagents.

Cells were routinely tested for mycoplasma contamination and negative results were recorded.

### Control of signaling level and duration

In experiments in which we controlled the level and duration of SMAD4 signaling at the population level, we used a modified experimental protocol. Cells were seeded in Ibidi μ-Slide 18-well plates at a low density of 1.5x10^4^ cells/cm^2^ 16-20 hours before treatment. To ensure uniformly high signaling, in addition to sparse seeding density, cells were maintained in RI over the course of differentiation and treated with a high BMP4 dose of 100 ng/mL. The signaling level was tuned by varying the concentration of BMPRi (LDN193189) added concurrently with BMP4, and duration was controlled by removing BMP4 and adding a saturating dose of 1000 nM BMPRi. Wnt ligand secretion inhibitor WNTi (IWP2) was included over the course of differentiation in all such experiments.

### Immunofluorescence staining

Samples were washed twice in PBS without calcium and magnesium (PBS^-/-^), fixed with 4% paraformaldehyde for 20 minutes at room temperature (RT), and then washed with PBS^-/-^ two more times. They were then incubated in a permeabilization buffer (0.1% Triton X-100 in 1X PBS^-/-^) for 10 minutes at RT and rinsed twice more with PBS^-/-^. Cell lines expressing fluorescent proteins were photobleached according to Lin et al., 2015 by incubating for 1 hour at room temperature in a bleaching buffer (3% H2O2, 20mM HCL diluted in PBS^-/-^) under an incandescent lamp, followed by two washes with PBS^-/-^. After permeabilization and optional bleaching, blocking was done with a blocking buffer (3% donkey serum + 0.1% Triton X-100 diluted in 1X PBS^-/-^) for 30 minutes at RT. Following blocking, samples were incubated overnight at 4°C in a solution of primary antibodies diluted in blocking buffer (antibodies and dilutions are listed in Table 2). Following primary antibody incubation, samples were washed 3x with PBST (0.1% Tween 20 in 1X PBS^-/-^), with 20 minutes incubation at RT between each wash. Samples were incubated in a solution of secondary antibodies diluted in blocking buffer (antibodies and dilutions described in Table 3) and DAPI (1 μg/mL; ThermoFisher Scientific) for 30 minutes at RT in the dark. After incubation, two fast PBST washes were performed, followed by two PBST washes with 20 minutes incubation at RT between each. Cells were stored in 1xPBS^-/-^ with 0.01% sodium azide, and were transferred to imaging buffer (700mM N-Acetyl-Cysteine in ddH2O, with pH adjusted to 7.4; Gut et al., 2018) immediately before imaging to prevent photocrosslinking of antibodies.

**Table 2:** Primary antibodies used for immunofluorescence.

| Protein | Species | Dilution | Catalog # | Vendor |
| --- | --- | --- | --- | --- |
| ISL1 | Mouse | 1:200 | 39.4D5 | DSHB |
| SOX2 | Rabbit | 1:200 | 3579S | Cell Signaling Technology |
| NANOG | Goat | 1:100 | AF1997 | R&D Systems |
| HAND1 | Goat | 1:200 | AF3168 | R&D Systems |
| GATA3 | Rabbit | 1:800 | 5852S | Cell Signaling Technology |
| TFAP2C | Mouse | 1:150 | SC-12762 | Santa Cruz Biotechnology |
| OCT3/4 | Mouse | 1:400 | 611202 | BD Biosciences |
| pSMAD1/5/9 | Rabbit | 1:100 | 13820S | Cell Signaling Technology |

**Table 3:** Secondary antibodies.

| Protein | Species | Dilution | Catalog # | Vendor |
| --- | --- | --- | --- | --- |
| Alexa Fluor 488 anti-mouse | Donkey IgG | 1:500 | A21202 | ThermoFisher Scientific |
| Alexa Fluor 555 anti-rabbit | Donkey IgG | 1:500 | A31572 | ThermoFisher Scientific |
| Alexa Fluor 647 anti-goat | Donkey IgG | 1:500 | A21447 | ThermoFisher Scientific |
| Alexa Fluor 647 anti-mouse | Donkey IgG | 1:500 | A31571 | ThermoFisher Scientific |

### Generation of the ESI017 tetO-SOX2 cell line

An enhanced piggyBac Puromycin selectable and DOX inducible vector was digested with EcoRI-NotI and ligated with PCR-amplified human SOX2 from the FUW-tetO-hSOX2 plasmid (Addgene#20724). Transfection into hPSCs was done using Lipofectamine Stem Reagent per the manufacturer’s instructions. Transfected cells were selected with 1ug/ml Puromycin for 1 to 2 weeks until only single colony clones remained.

### Repeat staining

To perform multiplexed immunofluorescence, we adapted the protocol described in Gut et al., 2018^41^ for elution and sample restaining. After the previous iteration of immunofluorescence imaging, the sample was washed three times with PBS^-/-^ and three rounds of antibody elution were performed; in each round the sample was incubated for ten minutes at room temperature in an elution buffer (0.5M L-Glycine, 3M urea, 3M guanidine hydrochloride, and 70mM TCEP, diluted in ddH2O, pH adjusted to 2.5) while being shaken at 50 rotations per minute (RPM) on a tabletop orbital shaker. The sample was washed three more times with PBS^-/-^, and blocked on the orbital shaker in blocking buffer for 30 minutes at room temperature. Primary antibody dilutions and incubation were done as for initial IF staining. Secondary antibody staining was performed as in initial staining, with the incubation in the solution of secondary antibodies extended from 30 min to an hour, and done on an orbital shaker at 50 RPM. Storage and imaging buffers are as for initial IF staining.

### Imaging

Imaging was performed with an Andor Dragonfly/Leica DMI8 spinning disk confocal microscope with a ×40, NA 1.1 water objective and a x20 air objective, as well as a Nikon/Yokogawa spinning disk confocal microscope with a x20 air objective. Live-cell imaging was performed with controlled temperature (37°), CO2 concentration (5%), and humidity (>60%). Experiments for which single-cell tracking was performed (fig 3) were performed using the x40 water objective. Other experiments were generally performed with a x20 air objective. Live-cell experiments in disordered culture were imaged every 10 minutes, using a z stack with 4 slices spaced 3 to 3.33 microns apart. Live-cell imaging of micropatterned colonies was done with a time interval of 20 minutes using a z stack with 4 slices spaced 4 to 5 microns apart. Media and treatment changes were performed in the time between imaging intervals without removing the sample from the microscope stage. For experiments in which both live and fixed-cell image data was quantified for the same cells, the same microscope and objective was always used; upon the conclusion of live-cell imaging, the sample was immediately taken for fixation to minimize the movement of cells between the conclusion of live-cell imaging and fixation and facilitate matching of live to fixed nuclei. PFA was added to the sample within 10 minutes of the conclusion of live imaging.

### Image analysis

In immunofluorescence data, nuclei were segmented based on DAPI staining using both Cellpose^62^ and the pixel classification workflow in Ilastik^39^. Ilastik and Cellpose masks were merged into a single segmentation as previously described^30^. After consolidating these segmentations, the object classification workflow in Ilastik was used to identify and discard missegmented (junk) objects. Cells in disordered culture form a monolayer and their nuclei were segmented based on the maximal intensity projection (MIP) of the z stack of DAPI images. For micropatterned colonies in which cells may be layered two or three cells deep, we segmented nuclei in each z slice and merged nuclear masks across z slices into a single 3D segmentation as previously described^30^.

In live imaging montages of micropatterned colonies, nuclei were segmented in the same way as in immunofluorescence data. In live cell images in sparser disordered culture, a nuclear segmentation pipeline optimized for single-cell tracking using only Ilastik was used. To facilitate single cell tracking after pixel classification, an additional step of Ilastik object classification marked segmented objects as interphase, metaphase (chromosomes aligned along the metaphase plate, immediately prior to splitting), other dividing (prophase -> chromatin is visibly condensed but not aligned; anaphase -> sister chromatids are moving apart but may not yet be segmented as two separate objects). For immunofluorescence data, an additional class was included to discard missegmented objects. Finally, a custom algorithm for approximate convex decomposition^30^ was applied to interphase-labeled foreground objects in the nuclear segmentation to split overlapping or touching nuclei into distinct masks.

Downstream quantification was carried out with a custom image-processing pipeline written in MATLAB. Expression levels were calculated as mean intensity in each channel within the nuclear mask. Both for segmentations of the MIP and 3D segmentations, the intensity was quantified for each nucleus in the z slice in which that nucleus was most in-focus, determined based on the intensity profile of the nuclear marker (DAPI or H2B) in z.

In live-cell images of GFP::SMAD4 or RFP:SMAD1, we additionally used Ilastik pixel classification to segment cell bodies as foreground, and used the inverse of this mask to detect the image background. To determine cytoplasmic intensity for each cell, a watershed operation was performed with nuclear masks imposed as minima for the watershed. For each nucleus, a cytoplasmic mask was constructed as an offset annulus about the nucleus, intersected with both the watershed basin corresponding to that nucleus and foreground mask of SMAD4 or SMAD1. Values in the cytoplasm were calculated as mean intensity in the cytoplasmic mask in the same z slice in which nuclear intensity was computed. Nuclear to cytoplasmic ratio was taken as background-subtracted nuclear intensity divided by background-subtracted cytoplasmic intensity.

For quantification of multiple rounds of immunofluorescence staining and imaging, phase correlation-based image registration was used to find a rigid shift aligning consecutive rounds of imaging to the first round. For cells in disordered culture, the same segmentation was used from the first round of imaging to quantify expression in subsequent rounds after alignment. For micropatterned colonies, a 3D segmentation was generated for each round of imaging separately, and individual cells were linked between rounds based on aligned x and y and normalized z centroid positions of each cell, using the algorithm for matching live to fixed cells described in the single-cell tracking supplement.

In micropatterned colonies of hPSCs, we performed analysis based on edge distance by subdividing the colony into 30 bins with equal numbers of cells in each bin. In each bin, all cells within that bin are within a similar distance from the colony edge. The average expression or signaling value for each bin was taken as the median among the cells in that bin.

### Single-cell tracking

Fully automated single-cell tracking was performed with a custom algorithm written in MATLAB described in detail in the supplemental text on the tracking algorithm, modified from Jaqaman et al^38^ and the implementation used in Trackmate^63^. Most importantly, to better handle cell division we applied machine learning^39^ to label nuclei as dividing based on morphological and image intensity information, and adjusted the linking function to handle nuclei during and after cell division (SI Fig. 2). For each single-cell tracking experiment, a subset of cell tracks were manually validated and results for the larger dataset were corroborated with the subset of validated tracks. Live cells were matched to fixed cells using the same algorithm used for tracking live cells as described in the supplemental text.

### Analysis of signaling histories

Analysis of single cell signaling histories was carried out in MATLAB and Python. In Fig. 1, clustering of signaling histories was done using soft k-means with k=3 in MATLAB with fcm. To compare the fate pattern to the signaling cluster pattern, we discretized the profile of fate markers, assigning the most prevalent fate at each position, and then averaged this over multiple colonies as a way to visualize (minimal) variation between colonies (Fig 1J, SI Fig 1G). The prediction of the fate boundary from signaling was initially less accurate with Wnt inhibitor than without (SI Fig. 1JK), but we found that this could be attributed to the fact that there is no objective way to assign the elbow fate, and the clustering algorithm produced the wrong assignment. Manually changing the cluster assignment of the elbow led to closer agreement with the fate pattern (SI Fig.1K).

In Fig. 3, signaling features were fit in MATLAB using lsqnonlin. Single cell histories were denoised in Python with MAGIC (Markov Affinity-based Graph Imputation of Cells) using three nearest neighbors (knn=3) and the diffusion operator to third power (t=3). After denoising the total variance explained of the leading three principal components went from 48% to 83% with all sub-leading components making very small contributions, suggesting that these sub-leading components mostly capture noise, consistent with the fact that they had no obvious interpretation (SI Fig. 3G).

Clustering by fate was first performed by fitting a two-component Gaussian mixture model to the seven-dimensional immunofluorescence data. We determined which markers best separate the clusters by calculating cluster separation as the difference in the means over to the sum of the standard deviations for a specific marker (SI Fig. 3JK). As an alternative approach, we also processed our seven-dimensional immunofluorescence data in the same way as single-cell RNA-sequencing data, clustered it with the Leiden algorithm and calculated differential expression between the clusters (SI Fig. 3LM). Both approaches yielded ISL1 and NANOG as top genes.

To determine the relationship between signaling and fate, we had the option of denoising both history and fate based on the cells with most similar fate marker expression, the cells with the most similar signaling histories, or some linear combination of the two. Therefore we compared the two extremes and to ensure we were not simply creating artificial correlations included a control where signaling histories were randomly assigned to marker expression before denoising, which reassuringly did not yield any correlation (SI Fig. 3L, bottom). We found that denoising based on fate yielded higher correlation between signaling and fate and better preserved the bimodal distribution of fate markers (SI Fig. 3O). Moreover, this approach is conceptually appealing because it directly extends the averaging over all cells with two discrete fates in Fig. 3H to essentially more fine-grained averaging of histories between small numbers of cells with most similar fate marker expression. We therefore applied fate-based denoising for combined analysis of signaling and fate.

To test how much information each feature contains about fate we determined the accuracy (% true positives + true negatives) of a Bayesian classifier, which is formally optimal^64^ and determines the most probable fate given the value of a signaling feature from the conditional probability P(fate|feature). Because of the monotonic relationship between fate and features this came down to determining an optimal threshold in the signaling feature. The four quadrants made by the fate threshold and the signaling threshold then provide the confusion matrix of the resulting binary classifier, with amnion-like cells (log(ISL1/NANOG) > 0) above/below the signaling feature threshold corresponding to true/false positive predictions, and pluripotent cells (log(ISL1/NANOG) < 0) above/below the signaling feature threshold corresponding to false/true negatives, respectively. From this one can then formally calculate the information contained about fate in the signaling features as a single number called the decoder-based mutual information^65^, which has the nice property that it is zero for pure chance, whereas the total accuracy (true positives + true negatives) is 50% for pure chance in this binary classification, but for simplicity we chose to present the accuracy.

### RNA sequencing and analysis

For RNA sequencing of the time series after BMP treatment, total RNA extraction was performed with the Invitrogran RNAqueous micro kit according to the manufacturer’s instructions. Cells were collected and lysed with the provided lysis buffer at specified times and the lysate was frozen and stored at -80°C. Whole RNA was prepared and DNase-treated for all samples at the same time, per kit instructions. The University of Michigan Advanced Genomics Core performed library preparation for mRNAs with ribosomal RNA depletion, and sequencing was performed in an Illumina NovaSeq S4 Flowcell with a sequencing depth of 57M reads per sample.

For the dose-response, total RNA extraction was performed with the QIAGEN RNeasy micro kit according to the manufacturer’s instructions. Five hours after BMP4 treatment cells were lysed with lysis buffer RLT and lysate was collected. Whole RNA was prepared and DNase-treated according to the kit instructions. The University of Michigan Advanced Genomics Core performed library preparation for mRNAs with polyA selection, and sequencing was performed in an Illumina NovaSeq S4 Flowcell with a sequencing depth of 33M reads per sample.

Reads from FASTQ files were trimmed using Cutadapt v2.3 (Martin, 2011) and mapped to the reference genome GRCh38 (ENSEMBL), using STAR v2.7.8a (Dobin et al., 2013). Count estimates were generated with RSEM v1.3.3 (Li and Dewey, 2011). Alignment options followed ENCODE standards for RNA-seq.

For analysis of time-series sequencing data, low-expressed genes were defined as those with less than 2.5 counts per million averaged over all conditions and filtered out. We further filtered out those showing relatively little change by keeping only genes with an absolute cumulative log2 fold change greater than ∼1.55 (one standard deviation). After filtering, each gene was normalized to its maximum value in the time series. Agglomerative hierarchical clustering was performed in MATLAB after normalization using euclidean distance and Ward’s linkage.

### Mathematical modeling

Dynamics of SOX2 and ISL1 expression were simulated with a nonlinear system of two first-order ordinary differential equations. Idealized SMAD4 dynamics with levels inferred from live imaging experiments were used as input to the model, and numerical simulations were performed in MATLAB. Rationale for the model construction and details of fitting to expression data are described in the supplemental text about the mathematical model.

## Supplementary Figures

**Figure 1 supplement:**
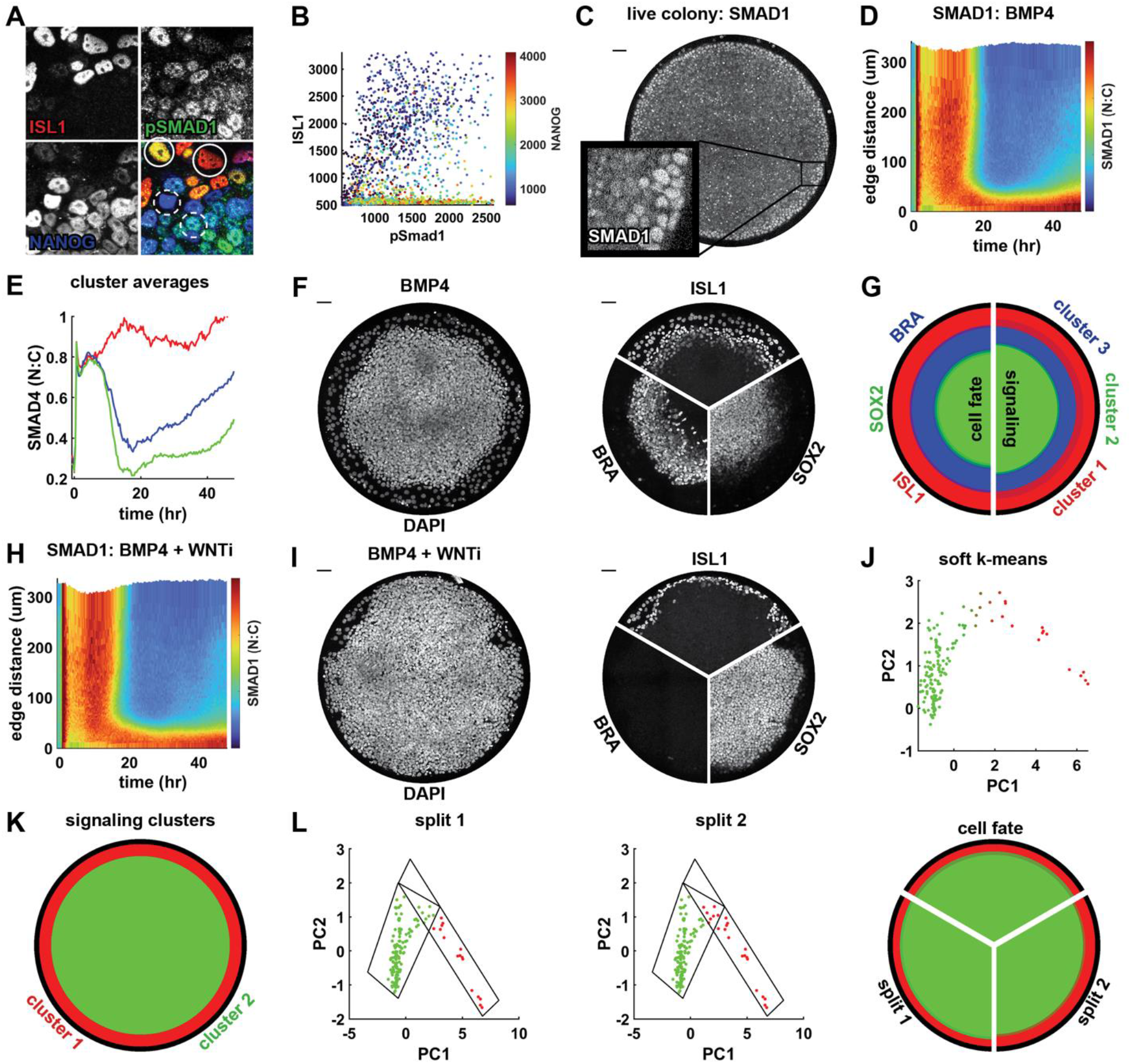
**(A-B)** detail of staining and quantification for pSMAD1, ISL1, and NANOG after BMP treatment in the presence of Wnt inhibitor (IWP2) showing low correlation between fate and final BMP signaling levels in a micropatterned colony. Solid circles indicate two amnion-like cells with high and low signaling levels respectively, and dashed circles similarly indicate two pluripotent cells at different signaling levels. (**C**) A representative micropatterned colony of RUES2 cells expressing RFP::SMAD1 at *t* = 30 hours after treatment with BMP4, showing nuclear localization of SMAD1 specifically at the colony edge. (**D**) Kymograph of mean SMAD1 signaling in N=5 micropatterned colonies treated with BMP4. (**E**) Mean signaling within the clusters of signaling histories in fig 1H. (**F**) Separated channel images showing the DAPI, ISL1, SOX2, and BRA stains corresponding to the colonies shown in Fig. 1J. (**G**) Comparison of the average profile of cell fate markers (left) and clusters of signaling histories (right) for BMP4-treated colonies. **(H)** kymograph showing average spatiotemporal dynamics of SMAD1 in micropatterned colonies treated with BMP4 and WNTi (IWP2). (**I**) Separated channel images showing the DAPI, ISL1, SOX2, and BRA stains corresponding to the colonies shown in Fig. 1N.**(J)** Scatterplot of the first two PCs of radially averaged signaling histories, colored for cluster assignment as in Fig 1G. **(K)** Predicted fate map based on the clustering in (J), determined as in Fig 1I. (**L**) Example of two different ways to assign the signaling histories corresponding to the ‘elbow’ of the PCA plot in colonies treated with BMP4 + WNTi, along with the resulting radial profiles for each assignment, compared to the profile of ISL1 and SOX2 expression in those colonies. Scale bars 50um.

**Figure 2 supplement:**
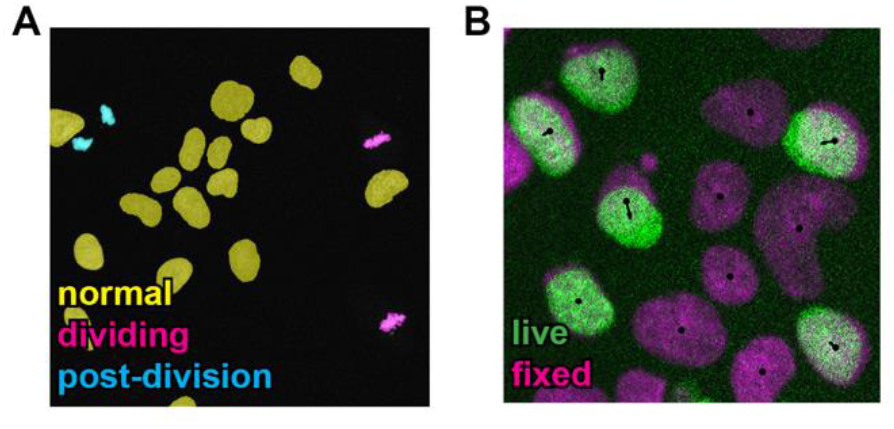
**(A)** Example image of nuclei with overlaid classification of cells as non-dividing (yellow), dividing (magenta), and immediately post-division (cyan). **(B)** Linking live to fixed cells. Image of live nuclei is shown in green, and fixed nuclei in magenta. Black arrows show links from the centroids of fixed nuclei to centroids of live nuclei.

**Figure 3 supplement:**
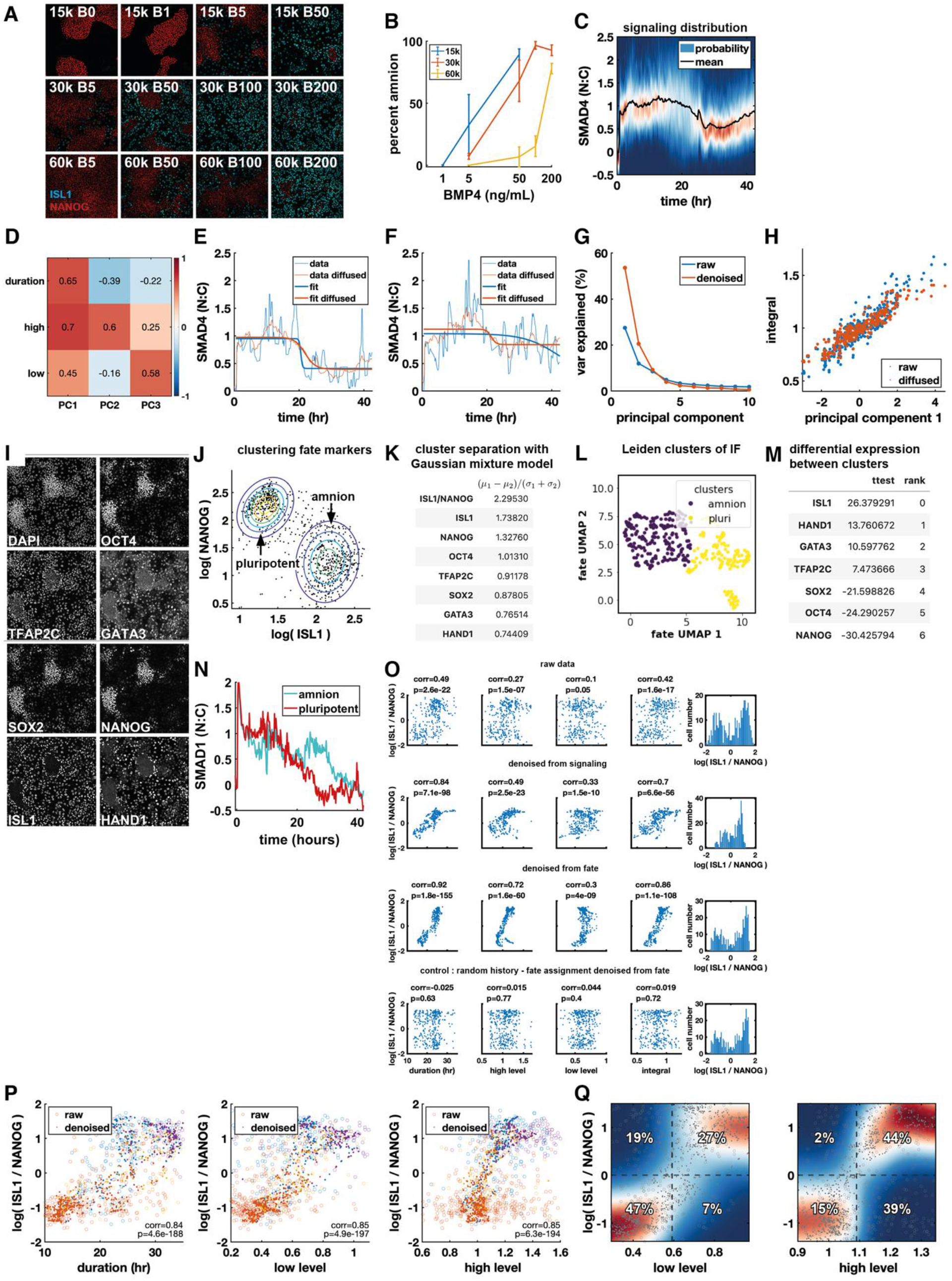
(**A**) immunofluorescence staining for NANOG and ISL1 for different conditions, first number is density, .e.g 15k = 15,000 cells / cm^2^, second number is BMP4 dose, e.g. B1 = 1ng/ml BMP4. **(B)** Quantification of (A). **(C)** Heatmap plot of signaling distribution over time corresponding to Fig. 3AB. **(D)** Correlation of features and principal components before denoising. **(E-F)** Example signaling histories before and after denoising via data diffusion with MAGIC, with sigmoid fits to the raw and denoised data. (**G**) Scatterplot of signal integral against principal component 1, with and without denoising. **(H)** Variance explained in signaling distribution from (C) by the first 10 PCs for raw and denoised signaling histories. (**I**) Representative single-channel IF images showing expression of all 7 stained genes in the same field of view. **(J)** Contour plot of a two-component Gaussian mixture model fit to fate marker expression, overlaid on a scatterplot of ISL1 vs. NANOG. (**K**) Table of values of a measure of cluster separation. The marginal distribution of the 7D Gaussian mixture model (GMM) is taken along each axis indicated and the separation of clusters along that direction is taken as the ratio of the difference in the means of the two GMM components to the sum of their standard deviations. A higher value indicates better separation. (**L**) UMAP plot showing the separation of cells into two clusters with Leiden clustering. (**M**) Table of differential expression of each marker between the Leiden clusters, showing highest absolute value for ISL1 and NANOG. (**N**) mean RFP::SMAD1 signaling in amnion and pluripotent cells. (**O**) Scatterplots of log(ISL1 / NANOG) and signaling features under various denoising schemes for data in Fig. 3A-M. **(P)** Scatter plots of signaling features vs. log(ISL1/NANOG) colored for condition with and without denoising for data in Fig. 3K-P. **(Q)** Heatmap of kernel density estimate after denoising of conditional distributions of log(ISL1 / NANOG) with respect to low level and high level of signaling, overlaid with a scatterplots of data points before (circles) and after denoising (dots). Dashed lines show separation of cells into amnion-like and pluripotent based on log(ISL1 / NANOG) or on signaling features. The percentage of cells in each quadrant is indicated, with correct assignments in the top right and bottom left quadrant of each heatmap.

**Figure 4 supplement:**
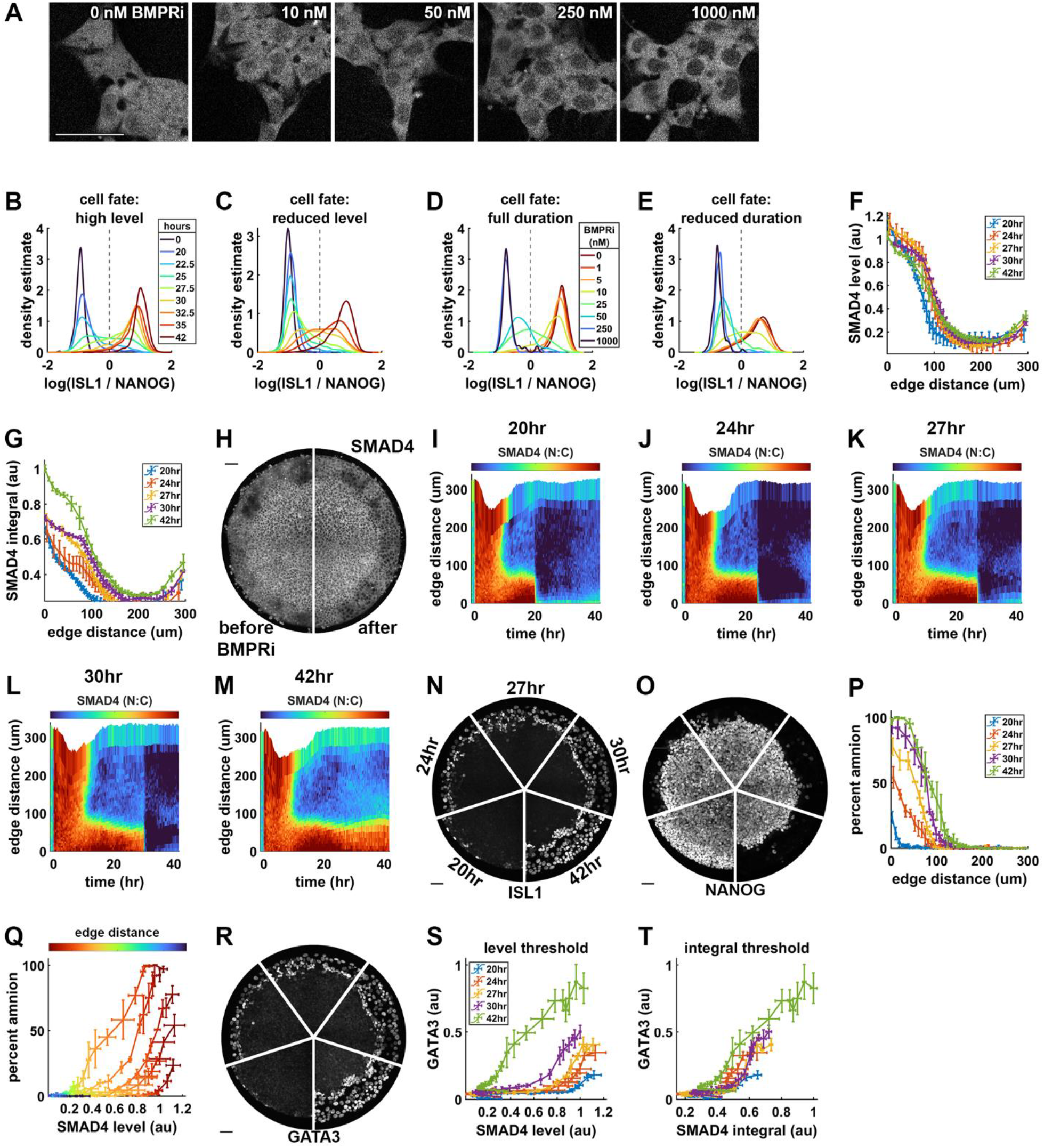
**(A)** GFP::SMAD4 images corresponding to data in Fig. 4H, showing sparsely seeded cells treated with 50ng/ml BMP4 and different doses of LDN193189 (BMPRi). (**B**) Kernel density estimate (KDE) of the log(ISL1/NANOG) distribution in each condition shown in 4D. (**C**) Kernel density estimate (KDE) of the log(ISL1/NANOG) distribution in each condition shown in 4E. Legend in (B). (**D**) KDE of the log(ISL1 / NANOG) distribution after 42h of differentiation for conditions in 4H. (**E**) KDE of the log(ISL1 / NANOG) distribution after 42h of differentiation for conditions in 4I. Legend in (D). **(F)** Average level of SMAD4 signaling before BMP inhibition as a function of distance from the colony edge for different durations of BMP signaling. **(G)** Integral of SMAD4 signaling at 42h as a function of distance from the colony edge for different durations of BMP signaling. (**H**) GFP::SMAD4 for a BMP treated micropatterned colony treated with 200ng/ml BMP4 shown before (29h, left) and after BMP signaling inhibition (31h, right). **(I-M)** Kymographs of average SMAD4 signaling in N=3 micropatterned colonies each for five signaling durations. **(N-O)** Representative IF images showing the spatial extent of ISL1 and NANOG expression in micropatterned colonies exposed to different durations of BMP signaling. **(P)** Quantification of percentage of differentiated cells (ISL1+NANOG-) as a function of distance from the colony edge for different durations of BMP signaling. **(Q)** Percent amnion differentiation vs. level of BMP signaling as in Fig. 4P, colored for distance from the colony edge. **(R)** Representative IF images showing the spatial extent of GATA3 expression in micropatterned colonies exposed to different durations of BMP signaling. **(S)** GATA3 expression in radial bins as a function of SMAD4 level before removal of BMP4 for each signal duration. Error bars are standard deviation over the same radial bin in N = 3 replicate colonies. **(T)** GATA3 expression in radial bins as a function of total SMAD4 integral. Error bars are as in E. Scale bars 50um.

**Figure 5 supplement:**
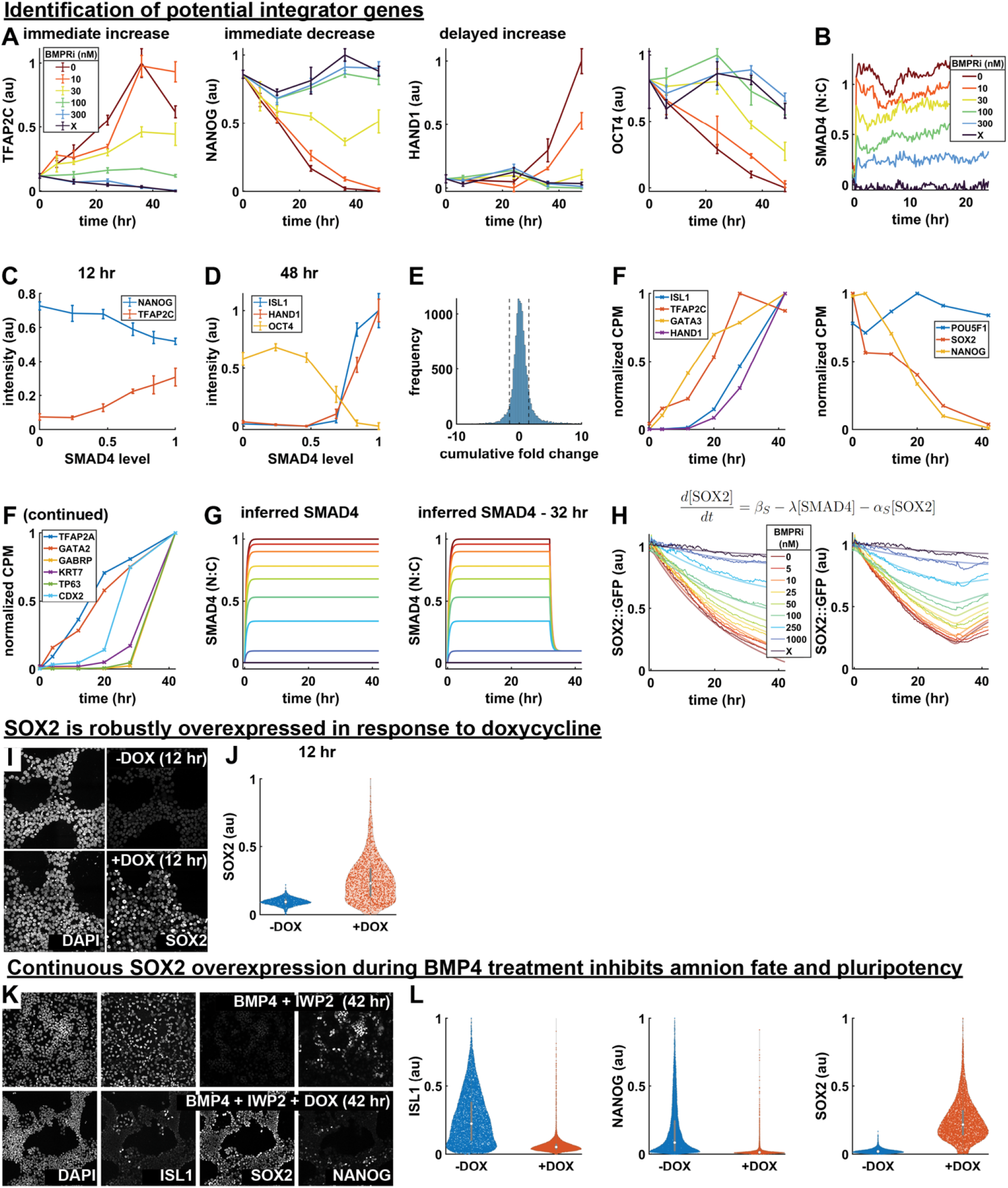
(**A**) Normalized expression of TFAP2C, NANOG, HAND1, and OCT4 over time for different signaling levels, measured with time-series IF. Error bars are standard deviation across N = 6 replicate images. (**B**) Average SMAD4 dynamics measured in the treatment conditions for which time-series IF was performed. (**C**) NANOG and TFAP2C expression at 12 hours, plotted against SMAD4 signaling level. (**D**) ISL1, HAND1, and OCT4 expression at 48 hours, plotted against SMAD4 signaling level, showing switch-like reliance. (**E**) Histogram of the cumulative log2 fold change over all genes in the time-series bulk RNA seq data. Genes with a fold change between the two dotted lines were not included in hierarchical clustering or subsequent analysis. (**F**) Expression over time for example genes measured with bulk RNA seq. Expression of amnion (left) and pluripotency (middle) genes that were also measured with time series IF, and additional amnion genes (right). (**G**) Idealized SMAD4 dynamics used as input to the ODE model, with level inferred from data in fig 4D. (**H**) GFP::SOX2 dynamics over the course of 42 hours of differentiation with indicated treatments applied for 42 (left) or 32 (right) hours, overlaid with fits of the simple ODE model described by the equation above it for SOX2, and the equation for ISL1 as in Fig. 4K. (**I**) Representative IF images showing expression of SOX2 and NANOG after 12 hours in pluripotency conditions with or without doxycycline. (**J**) Violin plot of SOX2 expression after 12 hours with or without doxycycline in pluripotency conditions. (**L**) Representative IF images showing ISL1, SOX2, and NANOG expression after 42 hours of treatment with BMP4 + WNTi with or without doxycycline. (**M**) Violin plots of ISL1, NANOG, and SOX2 expression with or without doxycycline in differentiation conditions.

## Algorithm for automated single-cell tracking

### Algorithm development

To construct tracks of single cells in time-lapse live-cell microscopy data, we took a “tracking by detection” approach (Magnusson et al. 2015), dividing the problem into two steps: (1) segmentation (detection) of all cells in each frame of the time-lapse, and (2) building tracks by linking segmented cells frame-to-frame. A custom single-cell tracking algorithm, based on the approach to particle tracking proposed in (Jaqaman et al. 2008), and similar to the implementation in the popular Fiji plugin Trackmate (Tinevez et al. 2017), was written in MATLAB and integrated into the image processing pipeline.

The approach taken in (Jaqaman et al. 2008) is to find an approximately optimal solution globally by breaking the tracking problem into two steps. Following this approach, we first link cells one-to-one or one-to-none in consecutive frames, assigning zero or one links from each cell in one frame to cells in the subsequent frame. This is followed by a “gap-closing, merging, splitting” (GMS) step, which addresses common segmentation and linking errors. Gap-closing connects the end of a track in frame *t*_1_ to the beginning of a track in frame *t*_2_ > *t*_1_ + 1, and is intended to account for nuclei leaving and re-entering the frame or that fail to be segmented in one or more frames. Merging connects the end of a track to the middle of another track, and accounts for two nuclei in frame *t* being segmented as a single nucleus in frame *t* + 1. Conversely, splitting connects the beginning of a track in frame *t* to the middle of a track in a previous frame, and accounts for two nuclei in frame *t* being segmented as a single nucleus in frame *t* − 1 or to a cell dividing in frame *t* − 1. These two steps are each cast as a linear assignment problem (LAP), in which a cost is assigned to each possible assignment and the globally optimal solution of the LAP minimizes the sum of possible costs. Fast algorithms have been developed to find the optimal solution for a given cost matrix, so the essential problem is to determine an effective way to assign costs to possible assignments, generally based on the proximity and morphological similarity of nuclei to be linked.

In addition to the general difficulty of robustly tracking through a time-lapse with segmentation errors, an additional challenge is tracking through cell division. To account for this, we modified the approach in Jaqaman to account for both segmentation errors and cell division. To facilitate the identification of dividing cells, we used the object-classification pipeline in Ilastik (Berg et al. 2019) to label all nuclei as dividing (M-phase, with chromosomes aligned along the metaphase plate immediately prior to cell division) or non-dividing. In the original algorithm, at the frame-frame linking stage each cell in frame *t* is linked to at most one cell in frame *t* + 1, and splits are only assigned later. We maintain this general framework, looking for only one daughter cell for each cell marked as dividing during frame-frame linking, and aiming to identify the second daughter cell at the merging, splitting, gap closing step. Additionally, the cost function for linking or splitting from

dividing nuclei are modified to facilitate identification of progeny cells, as described below.

In addition to modifying the frame-frame linking and GMS steps to better handle cell divisions, we add a step for merge resolution. This is motivated by the observation that merging events occur purely due to segmentation errors and so our final tracks should not incorporate the merging of two cells into a single object. To resolve merges, we first look for a split from the merged track, indicating that two nuclei moved close together and then apart again, and determine which of the input tracks to the merge more closely matches each of the output tracks from the split. If there is no subsequent split, we assume that either a merge was followed by separation of the two nuclei that failed to be detected as a split, or that the merge was assigned in error. In either case, we determine which input track to the merge more closely matches the track after the merge, and discard the other link.

### Frame-frame linking

To link cells in consecutive frames, we define the pairwise linking cost between each cell in frame *t* and each cell in frame *t* + 1, as well as the cost for ‘disappearance’ of cells from frame *t* and ‘appearance’ of cells in frame *t* + 1; that is, the cost for a cell in one frame to fail to be linked to any cell in the other. The cost for linking cell *i* in frame *t* to cell j in frame *t* + 1 is based on the cells’ xy positions, as well as the areas and intensities of the nuclei, and whether cell *i* is marked as dividing. The base cost for linking two cells is the squared euclidean distance between them, given by

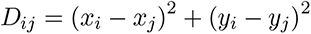

To obtain the final cost, we multiply this distance by weights based on the similarity of the two nuclei in area A and intensity I. If we define

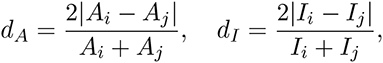

the the final cost is given by

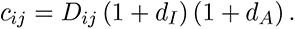

If cell *i* is labeled as dividing, an additional multiplicative weight is calculated based on the stereotypical rapid movement of daughter nuclei in opposite directions orthogonal to the orientation of the metaphase plate. This weight favors linking to prospective daughter cells found in a direction orthogonal to the metaphase plate. During image processing, the major and minor axes and orientation of an ellipse approximating the nucleus are calculated for each cell, and we use the orientation of cell i’s major axis as the orientation of the metaphase plate. We define a normalized vector *v̂* orthogonal to that orientation. We additionally define the vector pointing from cell *i* to cell j,

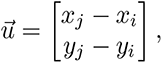

and normalize it to 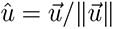. The additional weight for linking cell *i* to cell j is then

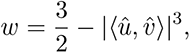

and the resulting overall cost is

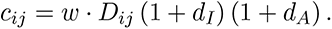

The inner product ⟨û, *v̂*⟩ depends on the angle between û and *v̂* and varies from −1 (antiparallel) to 0 (orthogonal) to 1 (parallel). Our weight then varies from 3/2 (orthogonal) to 1/2 (either parallel or antiparallel as daughter cells travel in both directions). Note that the range of values taken by w is unaffected by cubing the inner product, but results in a wider range of angles close to π/2 producing close to the maximum weight.

We may then construct the cost matrix A with rows corresponding to prospective links from the *n_t_* cells in frame *t* and columns corresponding to prospective links to the *n_t_*_+1_ cells in frame *t* + 1, so that A(i, j) = c*_ij_*, i.e.,

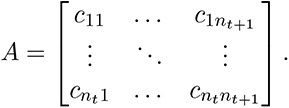

For computational efficiency, we additionally take as an input a maximum linking distance that defines the maximum distance a cell is expected to move between consecutive frames. We treat links between cells at a distance greater than this cutoff as impossible by setting the linking cost to Inf (arbitrarily large). In practice, we used a maximum linking distance of about 15 µm. We additionally define the alternative costs for appearance and disappearance for each cell to be 105% of the maximum finite linking cost. Cost matrices for link rejection are constructed as follows: *B*_1_ is an *n_t_* × *n_t_* diagonal matrix, with the cost for no link to be made to cell *i* in frame *t* at entry *B*_1_(*i*, *i*). All off-diagonal entries are set to Inf. Likewise, *B*_2_ is an *n_t_*_+1_ × *n_t_*_+1_ diagonal matrix storing the costs to reject links to cells in frame *t* + 1 and off-diagonal costs set to Inf. The resulting overall cost matrix is constructed as a block matrix as:

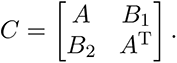

Assignments are made by choosing one cost in each row such that no two costs come from the same column and the sum of the costs is minimized. This optimization is performed with the Jonker-Volgenant algorithm for LAPs (Jonker and Volgenant 1987) implemented in MATLAB (Cao 2023). Note that the inclusion of the transpose of A in the lower corner ensures that the number of assignments is the same along the rows and columns so that C is a square matrix, and column indices of the assignments in the first *n_t_* rows will match the row indices of the last *n_t_* columns. Likewise, row indices of the assignments to the first *n_t_*_+1_ columns will match the column indices of the last *n_t_* rows.

### Gap closing, merging, splitting

The GMS step aims to tie up loose ends (and beginnings) from the frame-frame linking step. Track ends are cells without a link to a cell in a subsequent frame and track beginnings are those without a link from a cell in a previous frame (note that these are not mutually exclusive: if a cell has no link in the previous or in the subsequent frame it is both the beginning and end of its own one-cell track). Unlike in frame-frame linking, this step is not local in time and optimizes over possible assignments in the entire time series at once. Each track end is matched to either a track beginning (gap closing), a mid-point of another track (merging), or is given no assignment (track termination). Conversely, each track start is matched to a track end (gap closing), a mid-point of another track (splitting), or is not linked (track initiation). The structure of the cost matrix constructed to handle these possible assignments is more complex, and is constructed as a block matrix as

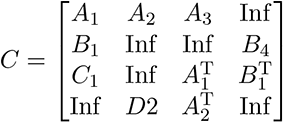

Here A_1_ contains costs for gap closing, A_2_ for merging, A_3_ for track termination, *B*_1_ for splitting, and C_1_ for track initiation. *B*_4_ is a diagonal matrix with costs to reject splits and D_2_ likewise has costs to reject merges. As in the frame-frame linking step, the cost matrix C is constructed to be a square matrix with the same possible assignments found along columns as along rows to satisfy the topological structure of the LAP. For instance, it can be seen that the first row of block matrices determines assignments from track ends, as does the third column of block matrices.

In constructing cost matrices, we again impose thresholds for computational efficiency, so links are only considered between cells within a maximum distance *δ*xy_max_ (in practice, about 22.5 um) and a maximum number of time steps apart *δt*_max_ (in practice, five frames).

The matrix A_1_ with costs for gap-closing is similar to the matrix of pairwise linking costs in the frame-frame linking step. Each entry of A_1_ stores the cost to link a track end at nucleus *i* in frame *t*_1_ to a track beginning at nucleus j in frame *t*_2_ with *t*_2_ > *t*_1_. If the nuclei are within the threshold distances of one another, *t*_2_ ≤ *t*_1_ + *δt*_max_ and ∥[*x_j_* − *x_i_*, *y_j_* − *y_i_*]*^T^*∥ ≤ *δxy*_max_, then the cost is given by

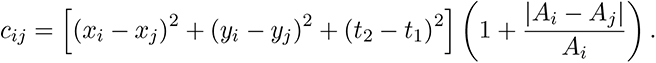

As in the frame-frame linking step, if cell *i* was labeled as dividing, this cost is multiplied by an additional weight w based on the angle between the normal vector to cell i’s major axis and the vector between cell *i* and cell j. Similar to the frame-frame linking step, we build a diagonal cost matrix A_3_ to reject links from each track end (cost for track termination) and C_1_ to reject links to each track start (cost for track initiation). Like in the frame-frame linking step, these costs are taken to be slightly larger than the maximum finite gap-closing cost.

The matrix A_2_ holds costs to merge track ends to midpoints of other tracks, where a track midpoint is any cell that is neither a track end nor a track start, i.e., that has a link both before and after it. Given cell *i* in frame *t*_1_ that is a track end, we find all track midpoints within the time and distance cutoffs of the track end. For a given midpoint cell j in frame *t*_2_ ≤ *t*_1_ + *δt*_max_, the cost to merge cell *i* into cell j is given by

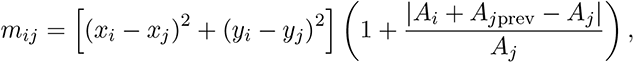

where A*_j_*_prev_ is the area of the nucleus preceding cell j in its track, so that the cost of accepting a merge is lowest when the area of the merged nucleus is the sum of the areas of the two input nuclei. The alternative cost matrix to reject merging holds the cost of rejecting merges for each midpoint for which at least one merge is considered, and is given by

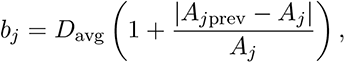

where D_avg_ is the averaged squared frame-frame displacement for tracks constructed in the frame-frame linking step. Then, the cost for rejecting a merge is lower than the cost of accepting the merge if |*A_j_*_prev_ − *A_j_*| < |*A_j_*_prev_ + *A_i_* − *A_j_*|, and if *D*_avg_ < (*x_i_* − *x_j_*)^2^ + (*y_i_* − *y_j_*)^2^.

The matrix *B*_1_ holds costs to split track starts from midpoints of other tracks. Similar to the construction of the cost matrix for merging, we take a track start cell *i* in frame *t*_1_, and find all track midpoints within the time and distance cutoffs. For a given midpoint cell j in frame *t*_2_ with *t*_1_ > *t*_2_ ≥ *t*_1_ − *δt*_max_ that is not marked as dividing, the cost to split cell *i* from cell j is given by

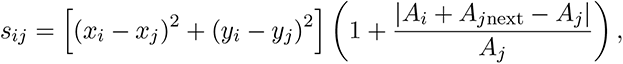

so the cost is lower if the area before the split is closer to the sum of the areas of the two cells after the split. If cell j is marked as dividing, however, we assume that the first link is to one daughter cell and attempt to find and link to the other daughter cell. The position of the first daughter cell is used to find the expected position of the other, based on the observation that immediately after division, sibling cells move symmetrically away from the location of the parent nucleus prior to division. To find the expected position of the remaining sibling nucleus, the displacement of the first sibling from the parent is found, and the expected position is taken to be at the same displacement but in the opposite direction. The linking cost then uses the distance of each prospective cell from this expected position instead of the distance from the parent cell itself. The resulting linking cost is

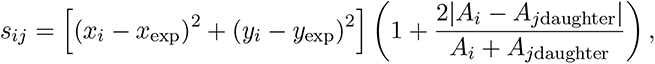

Where *A_j_*_daughter_ is the area of the first daughter nucleus in the same frame as the track start. Much like for merging, the alternative cost to reject splits is

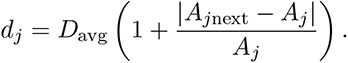

After all finite costs have been computed and the aggregate cost matrix is constructed as a block matrix, we numerically optimize to find the best solution to the LAP.

### Merge resolution

We resolve merges with the aim of separating the individual tracks that were inputs into the merge at later time points. If a split occurs from the same track soon after a merge (within the maximum time cutoff for gap-closing), and the nucleus from which the split occurred was not labeled as dividing, we assign each input to the merge to one output from the split and discard the links in between; otherwise, one of the links in to the merge is discarded, depending on which input cell bears greater morphological similarity to the cell after the merge. These one or two assignments to resolve each merge are made as a (very small) LAP with costs as in the frame-frame linking step.

### Linking live to fixed cells

At the end of live-cell imaging, each sample was fixed and immunofluorescence stained, and we include an additional step to link live cells at the end of the time-series to fixed cells. To ensure a consistent frame of reference, the image of nuclei in the last live frame is aligned to the DAPI stain of the fixed cells with phase correlation-based image registration and the positions of fixed nuclei are adjusted accordingly. Linking of individual cells is done in the same way as the frame-frame linking step during tracking, but with the linking cost based only on distance between nuclei and similarity in area. We do not consider nuclear intensity, which does not necessarily correlate between live data in which nuclei are labeled with fluorescent fusion proteins and fixed data where they are stained with DAPI. Sparse labeling introduces a potential complication to this step: only 10-20% of cells have nuclear markers in the live data, but every cell is stained for DAPI, including those that were not labeled live, so each live nucleus has potentially many more fixed nuclei nearby as candidates to which to link. However, we find that because there is little cell movement in the short time between the end of the live time lapse and the time that cells were fixed, our rigid image registration is robust to only a subset of nuclei being visible in the live data, and the alignment results in corresponding nuclei being very close in the aligned live and fixed data. To prevent erroneous linking to another nearby nucleus in the case that the true fixed nucleus corresponding to a given live cell failed to be properly segmented, we use a reduced maximum linking distance of 10 µm, or about one cell diameter, at this step.

### Additional implementation details

The algorithm was implemented in MATLAB, and incorporated into a larger image-processing pipeline.

The the optimal set of assignments for each LAP in the tracking pipeline is computed numerically with a MATLAB implementation (Cao 2023) of the Jonker-Volgenant algorithm (Jonker and Volgenant 1987).

To account for shifts in the entire field of view between consecutive time points, we implemented a “dejittering” algorithm. We iterated over all frames in the time lapse and at each time loaded a maximal intensity projection of the z stack of images of nuclei at time t*_i_* and t*_i_*_+1_ and used phase correlation to determine a global shift between the two images. At each time point, we applied the cumulative shift up to that time to the segmented cell positions in that time, effectively aligning the entire time-lapse to the field of view of the first frame. These updated cell positions are then used in the construction of cost functions for linking during tracking.

Parallelization is used to speed up the algorithm: at the frame-frame linking step, pairs of frames are linked in parallel, and in the gap closing, merging, splitting step, costs are computed in parallel as each block of the cost matrix is constructed.

## Mathematical model of BMP-SMAD4 integration

### Model development

We aimed to develop a mathematical model to explain how a simple gene regulatory network (GRN) could integrate BMP-SMAD4 signaling in time. In the simplest model, the expression of an integrator gene directly reflects the time integral of SMAD4 signaling. This is analogous to looking for genes with a rate of change that is a linear function of BMP-SMAD4 signaling, i.e., those that can be modeled with an ordinary differential equation (ODE) approximately as

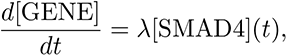

where [GENE] gives the concentration of protein and [SMAD4](t) is the (time-varying) level of BMP signaling. Integration of the above results in production of the gene product that is directly proportional to the signaling integral, i.e.,

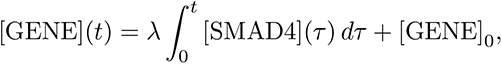

where the initial concentration of protein is given by [GENE]_0_ = [GENE](0). More generally, we can allow an additional constant term *β* for constitutive production, so that protein production remains a linear function of signaling. Additionally, as protein products are not indefinitely stable, we add a decay term that is proportional to the current concentration of protein. The ODE model then becomes

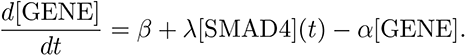

Our screen for genes for which the rate of change is linear with SMAD4 signaling level found SOX2 to be a promising candidate, as measured at the protein level with immunofluorescence, and at the transcript level with RNA sequencing. If SOX2 is our integrator, its dynamics should roughly follow integrated SMAD4 signaling, and it should repress late-response amnion genes so that they are expressed only if SOX2 goes below a threshold level. We first aim to determine whether SOX2 dynamics can be plausibly modeled with the dynamics described above. SOX2 is negatively regulated by BMP signaling, so we rewrite the ODE as

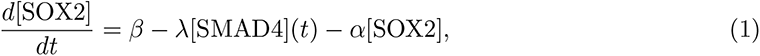

where *λ* is taken to be positive. In the absence of BMP signaling, [SMAD4] = 0 and SOX2 expression tends to a steady-state value of **β*/*α** balancing constitutive production and decay. A model of the form

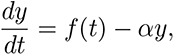

where *f* is a function of *t* but not y has the solution

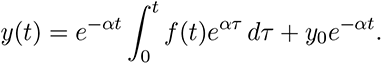

So the concentration of SOX2 is modeled by [SOX2](t) = *β*/*α* + ([SOX2]_0_ − *β*/*α*)e

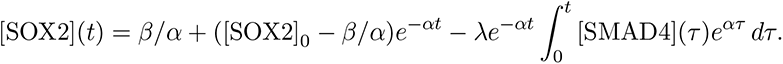

We assume that SOX2 is at or very near the steady state *β*/*α* in pluripotency maintenance conditions prior to stimulation of BMP signaling, i.e. [SOX2]_0_ = *β*/*α*. We further normalize this pretreatment expression level to one, so the expression for SOX2 as a function of time reduces to

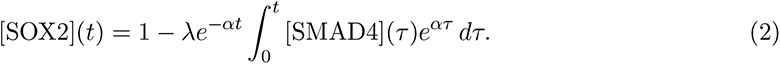

We see that the level of SOX2 protein reflects an exponentially-weighted integral of SMAD4 signaling, which closely approximates a direct integral of SMAD4 signaling for small *α* (where *e^−αt^* ≈ *e^αt^* ≈ 1). As a further simplification, if we consider conditions in which SMAD4 signaling is maintained at a steady level we can find the integral of [SMAD4] analytically and see that the SOX2 level exponentially decays towards the new steady state 1 − *λ* [SMAD4]/ *α* as described by

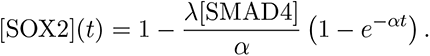

### Model fitting

To fit the model to experimental data, we measured GFP::SOX2 dynamics for 42 hours of BMP4-driven differentiation in conditions in which we could control the level and duration of signaling. Briefly, as described in the main text, we treated sparsely seeded hPSCs with a high dose of BMP4 to ensure uniformly high response, and controlled the signaling level via titration of a BMP receptor inhibitor (BMPRi). To control the duration, we shut down signaling by removing BMP4 and adding a high dose of BMPRi. We measured GFP::SOX2 dynamics at a range of signaling levels with durations of 42 and 32 hours (SI Fig 5H). As an indicator of differentiation to amnion, we measured ISL1 expression at the end of 42 hours with immunofluorescence. We approximated input SMAD4 dynamics as flat with levels measured in cells expressing GFP::SMAD4 in the same treatment conditions as the GFP::SOX2 cells (Fig 4; SI Fig 5G). We confirmed the linear relationship between SMAD4 signaling level and rate of SOX2 decay with a linear fit to the first 16 hours of GFP::SOX2 dynamics in each condition (Fig 5H). To determine the values of the parameters *α*, *β*, and *λ* in the model, we collected values of SOX2, SMAD4, and the slope of SOX2, averaged over short time windows to reduce the effect of measurement noise, and fit a plane defined by

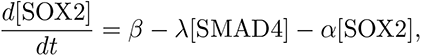

to our measured values of d[SOX2]/dt, [SOX2], and [SMAD4]. Numerically integrating the model with the fitted parameters and with measured input SMAD4 dynamics, we see generally good agreement with measured SOX2, but with some discrepancies between model and data appearing later in the course of differentiation in conditions with the highest levels of signaling (SI Fig 5H, red and orange curves). In particular, the model predicts faster initial decay followed by a plateau at later times, and for conditions in which BMP signals are removed, the model predicts sharp recovery of SOX2 levels. Signaling is completely shut down after addition of a high dose of BMPRi, so SOX2 should exponentially approach the initial level in all of these conditions. Because of this, the initial slope of recovery is expected to be highest for those with the lowest SOX2 level at the time of BMPRi addition. In contrast, SOX2 fails to robustly recover upon addition of BMPRi in high-signaling conditions (SI Fig 5H, right). Notably, there is a more pronounced recovery in SOX2 levels after BMPRi addition in conditions with lower signaling. We hypothesized that failure of SOX2 to recover after signaling shutdown in conditions with higher initial signaling reflected commitment to exit pluripotency. Because this effect seems to be pronounced only at later times in high-signaling conditions, we took it to be caused by repression of SOX2 by late-response amnion genes which only turn on in those conditions, and used ISL1 as a representative example of that class of genes. We modeled SOX2 as repressing expression of ISL1, as expected if it is our integrator gene. We additionally modeled repression of SOX2 by ISL1 so that SOX2 expression is further downregulated once ISL1 begins to be expressed. This fits the paradigm of mutually inhibitory regulatory programs specifying distinct cell fates that are widespread in development (Levine and Davidson 2005, Deĺas and Briscoe 2020). To implement this mutual repression mathematically, we take each gene to act on the other with Hill function dependence:

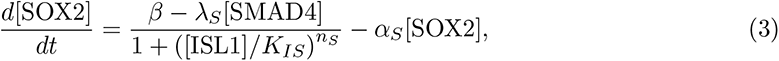

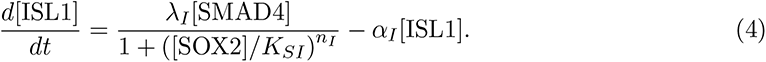

In the above equations, the parameters K describe the threshold for 50% inhibition of one gene by the other and n describes the steepness of the Hill function. In our time series expression data for ISL1 (Fig 5A, SI Fig 5F), we see that it remains close to zero until 20-24 hours, when expression rapidly switches on, suggesting that there is a sharp threshold for regulation of ISL1. We therefore modeled repression of ISL1 by SOX2 with a switch-like Hill function by setting *n_I_* = 4. On the other hand, repression of SOX2 by the amnion transcriptional program appears more graded, and we take *n_S_* = 2. We used simulated annealing to fit equations (3) and (4) to measured SOX2 and ISL1 expression data, using the same SMAD4 input dynamics described above. Briefly, the Values of each parameter must be initialized: we used values found with the previous fit of SOX2 alone, i.e., *λ _I_* = *λ _S_* = *λ*, *α _I_* = *α_S_* = α, and *β_S_* = *β*. We further initialized the inhibition threshold coefficients K*_SI_* and K*_IS_* at 0.5. We then numerically evaluated the system of ODEs (3) and (4) with those parameters and the SMAD4 inputs described above, and calculated the mean squared error E between the target and calculated expression levels. Then for a set number of iterations, we do the following: perturb the parameters by applying Gaussian noise to each with a variance of 10*^−^*^5^ and calculate the new error E_new_ after running the model with the new parameters. If the new error is lower, accept these values as the new parameter values; otherwise, we may still accept the new parameter values with probability exp (−Δ*E/k_B_*T), where Δ*E* = *E*_new_ − *E*, *t* is the ‘effective temperature’ for the annealing, and k*_B_* is a tunable constant. The value of *t* linearly decreases to zero over the course of the iterations so that accepting a set of parameters resulting in a higher cost becomes increasingly unlikely as the algorithm progresses, allowing exploration of the parameter space at early iterations to avoid becoming trapped at a local minimum in the parameter landscape, and settling in to a specific minimum at the end.

We see that the resulting simulated SOX2 dynamics align more closely with the measured dynamics, and resolve the discrepancies mentioned above (Fig 5I). Furthermore, we see that the relationship between the SMAD4 integral and ISL1 expression seen in the data is conserved for both signaling durations in our model (Fig 5J).

To generate the results shown in Fig 5IJ, we used the following parameter values:

| Parameter | Value | Meaning |
| --- | --- | --- |
| $(\alpha_S, \alpha_I)$ | (0.0363, 0.09) | protein dilution + degradation rates |
| $\beta$ | 0.0329 | constitutive SOX2 production rate |
| $(\lambda_S, \lambda_I)$ | (0.0397, 0.1240) | coefficients for regulation by SMAD4 |
| $(n_S, n_I)$ | (2, 4) | Hill function coefficients |
| $(K_{SI}, K_{IS})$ | (0.266, 0.3995) | inhibition thresholds |

## Notes

### Competing Interest Statement

The authors have declared no competing interest.

